# The Silent Saboteur: How Mitochondria Shape the Long-Term Fate of the Injured Brain

**DOI:** 10.1101/2025.03.19.644244

**Authors:** Sahan BS Kansakar, Sydney P Sterben, Charitha C Anamala, Mitchell D Thielen, Volha Liaudanskaya

## Abstract

Traumatic brain injury (TBI) is a major risk factor for neurodegenerative diseases, including Alzheimer’s disease (AD), yet the mechanistic link remains unclear. Here, we integrated human patient-derived transcriptomics with a 3D in vitro brain injury model to dissect cell-specific mitochondrial dysfunction as a driver of injury-induced neurodegeneration. Comparative transcriptomic analysis at 6 and 48 hours post-injury revealed conserved mitochondrial impairments across excitatory neurons, interneurons, astrocytes, and microglia. Using a novel cell-specific mitochondria tracking system, we demonstrate prolonged neuronal mitochondrial fragmentation, bioenergetic failure, and metabolic instability, coinciding with the emergence of AD markers, including pTau, APP, and Aβ42/40 dysregulation. Glial mitochondria exhibited delayed but distinct metabolic dysfunctions, with astrocytes impaired metabolic support and microglia sustained chronic inflammation. These findings establish neuronal mitochondrial failure as an early trigger of injury-induced neurodegeneration, reinforcing mitochondrial dysfunction as a therapeutic target for preventing TBI-driven AD pathology.

## Introduction

Traumatic Brain Injury (TBI) affects an estimated 69 million people annually ^1^. TBI is a debilitating condition with multifaceted pathobiology that leads to long-lasting disabilities and morbidity. What begins as a mechanical injury quickly launches a complex cascade of poorly understood molecular and metabolic events, with intercellular interactions amplifying and progressing the injury ^2,3^. Although the primary insult may produce significant brain damage, the secondary injury cascades — including mitochondrial dysfunction, metabolic alterations, neuroinflammation, axonal degeneration, persistent inflammation, protein misfolding (alpha-synuclein (aS), amyloid precursor protein (APP), tubulin-associated unit (Tau), transactive response DNA-binding protein of 43 (TDP43)), and cell death — play a critical role in injury outcomes. Chronic TBI pathology varies with patients, where some recover completely while others suffer physical and cognitive decline, leading to the onset of neurodegenerative diseases ^4^. Numerous epidemiological studies have demonstrated that TBI is the third strongest risk group for individuals to develop AD and AD-related dementias (ADRD) ^5–7^ after age and APOE genotype ^8^. While TBI-associated clinicopathological studies have demonstrated that amyloid beta (Ab) plaques and neurofibrillary tangles (NFTs) consisting of hyperphosphorylated Tau (p-tau) significantly contribute to AD/ADRD development, mechanisms are still poorly understood ^8,9^.

Mitochondria are a potential regulator of secondary injury progression and injury-induced AD-like neurodegeneration^10^; their dysfunction is a common feature of various diseases, including neurodegeneration and aging, metabolic disorders, and cancer. Disruption in mitochondria function influences inflammatory pathway activation, energy production, cellular metabolism, calcium regulation, transcription factor phosphorylation, and cell death activation ^11,12^. Various *in vivo, in vitro,* and human athletes studies demonstrated that TBI disrupts mitochondria function and metabolism (TCA cycle-related enzymes ^13–15^, mitochondria metabolism ^16–20^, fructose metabolism ^21^, and lipid peroxidation ^22,23^), implicating these alterations with secondary neurodegeneration and neuroinflammation ^24,25^. In addition, studies linked mitochondrial dysfunction — particularly impaired mitochondrial respiration — and decreased adenosine triphosphate (ATP) production to oxidative damage of neuronal lipids, nucleic acids, and proteins ^26^, accumulation of Aβ and p-Tau proteins ^8,9^, and development of AD ^27,28^.

Despite overwhelming proof of injury-induced mitochondria dysfunction and its association with an AD-like neurodegenerative phenotype, some critical gaps remain: Does mitochondrial dysfunction vary across cell types? Can mitochondria recover over time post-injury? Is there a correlation between cell-specific mitochondria dysfunction and AD-like neurodegenerative progression? However, most importantly, why are there still no answers to these questions? Although human studies are invaluable in providing information on the brain response to trauma, they struggle with causal validation of the molecular mechanisms associated with injury progression. Similarly, rodent TBI models present challenges in interrogating cell-specific contribution and molecular footprint due to the differences in genetic background, complexity, and pricing of such studies ^30^. Thus, there is a critical need for an alternative approach using human data to build a relevant hypothesis for validating cell-specific molecular mechanisms that drive the progression of brain injury-induced neurodegeneration and neuroinflammation.

Based on human patients’ data and extensive literature review, we hypothesize that injury-induced dysregulation in neuronal mitochondria bioenergetic and metabolic functions, with glial cells failing to provide long-term support, transform them into neurodegenerative factories that control the onset of acute neuroinflammation and chronic neurodegeneration, similar to AD phenotype. To address this hypothesis, we have integrated fluorescent tags with different wavelengths in brain cells (neurons (dsRED2), astrocytes (EGFP), and microglia (BFP2)) under mitochondria-specific promoter COX8A (cytochrome c Oxidase subunit 8A). These novel lines were used to triculture in the human 3D brain-like injury model ^29–33^, to unveil cell-specific alterations of mitochondria and their contribution to AD-like neurodegeneration progression post-TBI. While organoids are exceptional in studying developmental or genetic disorders, bioengineered 3D *in vitro* models might be a better fit to investigate injury-associated molecular alterations as they provide mechanical strength to withstand the injury, allow manipulation of cellular composition, and can have a better resemblance of extracellular structure^34,35^.

This study provides a comprehensive, cell-specific characterization of mitochondrial dysfunction following traumatic brain injury (TBI) and its potential role in driving Alzheimer’s-like neurodegeneration. We first validated acute transcriptomic alterations by expanding our initial 24-hour analysis to 6 and 48 hours, revealing early mitochondrial stress responses, transient compensatory mechanisms, and subsequent metabolic shifts. To assess whether these alterations translate to human-relevant neurodegenerative processes, we identified cell-specific transcriptomic convergence between our 3D *in vitro* human brain injury model and TBI patient-derived datasets ^36^, demonstrating that mitochondrial impairments in neurons, astrocytes, and microglia align with those observed in human pathology. To further dissect mitochondrial dysfunction at a cellular level, we developed and validated brain cell lines with fluorescently labeled mitochondria to precisely visualize and functionally assess mitochondria post-injury. Using this system, we uncovered distinct mitochondrial dysfunction trajectories across cell types, with neuronal mitochondria exhibiting persistent fragmentation and bioenergetic failure, astrocytic mitochondria demonstrating delayed metabolic collapse, and microglial mitochondria expressing chronic inflammatory activity. We correlated these mitochondrial impairments with the progressive accumulation of AD-related markers, including pTau, APP, and Aβ42/40 dysregulation, reinforcing the idea that mitochondrial failure serves as an early driver of TBI-induced neurodegeneration. This study is the first of its kind to integrate cell-specific transcriptomics, live mitochondrial imaging, and bioenergetic and metabolic profiling in a human-relevant TBI model, providing unprecedented insight into how mitochondrial dysfunction may initiate and sustain long-term neurodegenerative processes.

## Results

### Acute transcriptomic alterations in mitochondria function in human 3D *in vitro* brain tissue model

A previous study ^30^ with a 24-hour transcriptomic dataset demonstrated significant alterations to molecular pathways associated with mitochondria bioenergetic and metabolic functions, specifically in Mitochondrial Complex I gene expression. Additionally, the significant upregulation of fatty acids oxidation, carnitine shuttle, and mitochondria central dogma genes was shown to be crucial in mitochondria maintenance and repair. While the original study was the foundation for providing the starting information on mitochondria alterations, it was limited to one-time points, thus not allowing for dynamic study. Here, we first performed an extended analysis of responses at 6 and 48 hours to complete the transcriptomic profile of acute mitochondria dysfunction.

We used the CEMiTool R package ^37,38^ to group differentially expressed genes (DEGs) into modules at 6 and 48 hours post-injury (Figure S1), identifying five distinct modules. One module (Module 4) did not show significant changes in response to trauma and was excluded from further analysis (Figure S1-S2). The remaining four modules displayed dynamic shifts in gene expression, indicating early metabolic and signaling alterations followed by activation of cell cycle regulation and inflammation (Figure 1 A-B).

**Figure 1.**
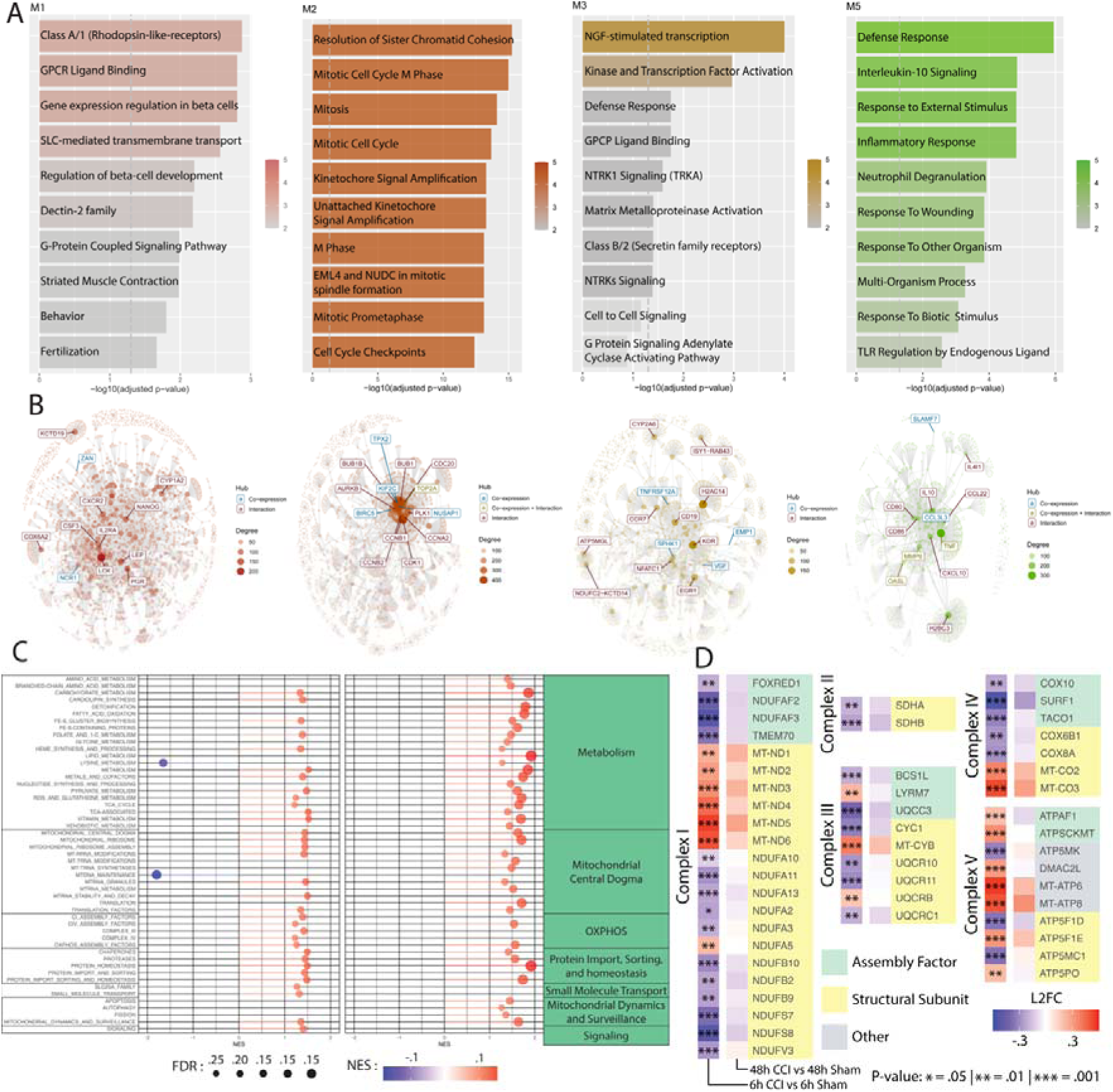
**Transcriptomic profile demonstrated acute mitochondrial damage** in human 3D *in vitro* brain tissues 6 and 48 hours after moderate injury inflicted by the controlled cortical impactor. The human 3D *in vitro* triculture model comprised 2 mln of neurons, 0.5 mln astrocytes, and 0.1 mln of HMC3 microglial cells. Sham (n = 5) and CCI (n = 6) at 6 hours, and Sham (n = 4) and CCI (n = 4) were used to obtain data for this transcriptomic profile. (A) Gene set enrichment analysis revealed 6 distinct expression modules associated with M1 – mitochondria function (positively associated with an injury); M2 – cell cycle progression (negatively associated at 6 hours); M3 – cellular signaling (positively associated with injury at 6 hours and negatively at 48 hours); M5 – neuroinflammation (negatively associated at 48 hours). (B) Critical regulators of molecular mechanisms associated with M1, M2, M3, and M5 modules. The full map of 6 detected modules and corresponding pathways can be found in Supplementary Figures S1 and S2 A-B. (C) Differentially expressed genes (DEGs) associated with mitochondria pathways and (D) Mitochondria electron transport chain complex genes were detected (abs(Log2(Fold Change))>0.585 and q<0.01).

Module 1, associated with the 6-hour time point, exhibited increased gene expression following injury, with COX6A2 (cytochrome c oxidase subunit 6A2) emerging as a key hub, directly linking this module to mitochondrial bioenergetic function. Module 2 showed a sharp decline in gene expression at 6 hours, primarily in genes related to cellular division, suggesting an early cell cycle arrest. However, by 48 hours, these genes were upregulated, indicating a shift toward cell proliferation and mitotic activity. Module 3 demonstrated a transient spike in transcriptional signaling pathways, including Nerve Growth Factor (NGF), Nuclear Factor kappa-light-chain-enhancer of activated B cells (NFkB), Neurotrophic Receptor Tyrosine Kinase 1 (NTRK1), and G Protein-Coupled Receptor (GPCR) -associated pathways, at 6 hours, with gene expression returning to baseline by 48 hours. Key mitochondrial hubs in this module, ATP5MGL (ATP Synthase Membrane Subunit g-Like) and NDUFC2-KCTD14 (a read-through fusion gene of NADH: Ubiquinone Oxidoreductase Subunit C2 and Potassium Channel Tetramerization Domain Containing 14), suggest a direct link between these early transcriptional responses and mitochondrial adenosine triphosphate (ATP) production. Lastly, module 5 displayed downregulated gene expression at 48 hours, was enriched for inflammatory response pathways, with hubs including CCL3L3 (C-C Motif Chemokine Ligand 3-Like 3), Tumor Necrosis Factor (TNF), CXCL10 (C-X-C Motif Chemokine Ligand 10), CD80 (Cluster of Differentiation 80), CD86 (Cluster of Differentiation 86), Interleukin 10 (IL-10), and CCL22 (C-C Motif Chemokine Ligand 22) highlighting the de-activation of immune signaling at later time points.

Next, we used Gene Set Enrichment (GSEA) to analyze the MitoCarta3.0 dataset (Figure 1 C-D) to evaluate enriched metabolic pathways acutely affected by injury ^39–42^. The overall cellular metabolic state was upregulated acutely after injury (including 6, 24, and 48 hours), alluding that injury induces hypermetabolism acutely after trauma, including pathways associated with OXPHOS, mitochondria maintenance and repair, metabolism and signaling. While overall cells demonstrated hypermetabolism, electron transport genes (ETC) at the acute 6-hour time point (6h CCI vs 6h Sham) demonstrated significant downregulation of Mitochondrial Complex I gene expression (Figure 1C), apart from the mitochondrially encoded NADH dehydrogenases (MT-ND) gene family (the gene family responsible for the onset of electron acceptance in the inner mitochondria membrane to initiate the proton gradient). Similar to Complex I, Complex II-IV also demonstrated downregulated genes. However, genes in Complex V were significantly upregulated. 48 hours after the injury, these transcriptomic alterations (48h CCI vs 48h Sham) return to baseline, with none of them significantly different. On the contrary, the 24-hour timepoint ^30^ showed a drastic increase in the complex I genes, with a few genes being differentially expressed in the other complexes. Unlike in 6 hours, complex V was almost unchanged.

### Multicellular mitochondria dysfunction as the convergent signature of brain trauma between human patients and human 3D *in vitro* model

In the previous section, we have shown that injury induces severe changes to mitochondrial function. In addition to our studies, there is a consensus in the field of brain trauma that mitochondria dysfunction is a potential regulator of injury-induced neurodegenerative alterations for disorders (Alzheimer’s, Parkinson’s, Chronic Traumatic Encephalopathy) ^5–7^.

While our study and many others suggest the involvement of mitochondria metabolism in injury-induced neurodegeneration, one question remains unknown: is this a general phenomenon, or does the mitochondria of different brain cells respond differently to brain trauma? To address whether human patients demonstrate cell-specific mitochondria dysfunction and how these responses differ, we focused on understanding cell-specific metabolic pathway alterations convergent between our bulk RNA-seq data (3D *in vitro* brain injury model) and single-nuclei RNA-seq (snRNA-seq) data (TBI human samples) ^36^. We first used bulk RNA-seq data from our 3D *in vitro* brain injury model to compare with the Reactome Pathway dataset, MitoCarta 3.0 dataset, and Mitochondrial Complex Gene dataset to identify altered pathways post-injury (Figure 2A and Figure S3A). We then used cell-specific human patient single-nuclei RNA-Seq data to identify how similar DEGs between these two samples contribute to altered pathways (Figure 2B and Figure S3B).

**Figure 2.**
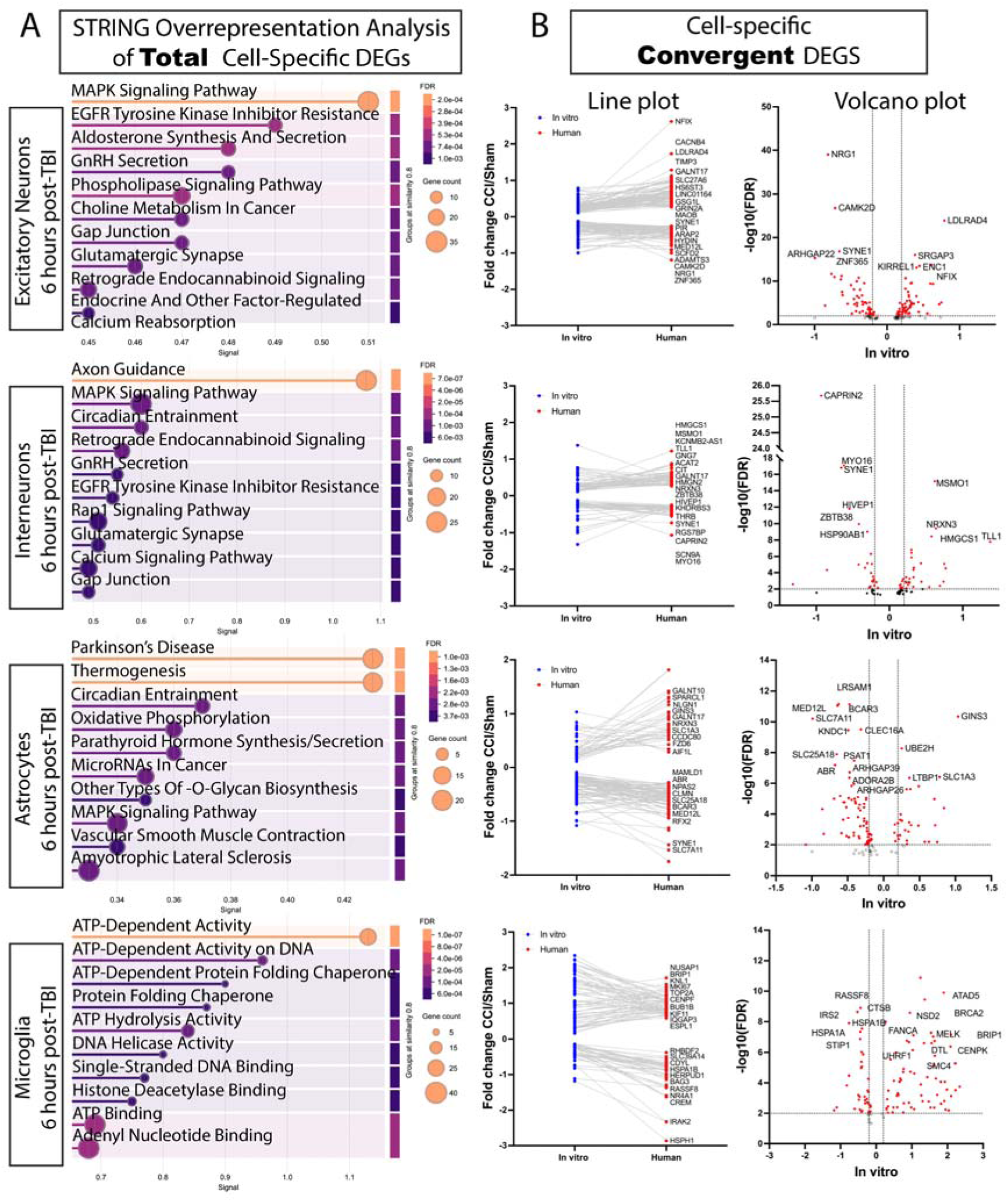
Human TBI transcriptomics align with our 3D in vitro model, revealing conserved mitochondrial dysfunction. The transcriptomic profile of human TBI patients^36^ revealed a cell-specific mitochondrial dysfunction signature that converges with our human 3D *in vitro* brain tissue model (comprised of 2 mln of neurons, 0.5 mln astrocytes, and 0.1 mln of HMC3 microglial cells) at 6 hours post-injury (48 hours results can be found in Supplementary Figure S3), highlighting conserved metabolic vulnerabilities across experimental and clinical contexts (n=5 Shams and n=6 CCI samples at 6 hours and n=5 Shams and n=6 CCI samples were used to obtain data for this transcriptomic profile). Single-cell RNA sequencing from 12 critically injured or deceased TBI patients was analyzed to curate cell-specific differentially expressed genes (DEGs) across excitatory neurons, interneurons, astrocytes, and microglia, using data from five uninjured controls as a reference. This patient-derived transcriptomic dataset was then integrated with bulk RNA sequencing from human 3D in vitro brain injury samples at 6 and 48 hours post-injury, enabling a comparative analysis of conserved molecular signatures between clinical and experimental models. Differentially expressed genes (abs(Log2(Fold Change))>0.585 and q<0.01). (A) Regardless of convergence or divergence, similar DEGs across the datasets were run with STRING analysis to determine cell-specific pathways significantly altered in response to trauma. (B) Convergent upregulated and downregulated DEGs were isolated and plotted for 4 distinct cell populations.

We used a publicly available snRNA-seq dataset that collected samples from 12 critically injured or deceased patients from brain trauma and curated cell-specific (excitatory neurons, interneurons, astrocytes, and microglia) differentially expressed genes (DEGs) compared to 5 control patients ^43^. Regardless of convergence or divergence, we selected similar DEGs across the datasets for STRING analysis to determine cell-specific pathways significantly altered in response to trauma (Figure 2A-B; Figure S3 A-B) 6 and 48 hours after injury. The results demonstrated pathways in excitatory neurons and interneurons were mostly associated with structural and synaptic network repair and electrical activity 6 and 48 hours post-injury. In addition, the transcriptomic profile also demonstrated dysfunction in Mitogen-Activated Protein Kinase (MAPK) signaling, critical pathway regulators of mitochondria function, oxidative stress, inflammation, and apoptosis.

While neuronal cells attempted to restore their function, both glial cells demonstrated drastic changes to ATP production via the oxidative phosphorylation (OXPHOS) pathway. At 6 hours post-injury, astrocytes demonstrated alterations in the MAPK pathway and O-glycan biosynthesis, and microglia demonstrated ATP production-related mechanisms and potential epigenetic alterations (Figure 2A). At 48 hours post-injury, astrocytes demonstrated changes to pyrimidine, purine, and nicotinamide metabolism, as well as microglia inflammation activation. Both glial cells demonstrate an association with axonal guidance (Figure S3 A-B).

Next, we focused on convergent DEGs between human patients and our human 3D *in vitro* brain model. At 6 hours, the analysis demonstrated alterations in Calcium/Calmodulin-Dependent Protein Kinase II Delta (CAMK2D) ^44–46^ and Monoamine Oxidase B (MAOB) ^47^ gene expression, with CAMK2D influencing synaptic plasticity and memory formation and MAOB contributing to oxidative stress and neurodegeneration through excessive production of hydrogen peroxide, both of which are disrupted in AD-phenotype. Though not directly connected to AD, some DEGs like CTSB (cathepsin B), Solute Carrier Family 7 Member 11 (SLC7A11) (glutathione metabolism) ^48^, and B-cell Lymphoma 6 (BCL6) (inflammation) may facilitate neurodegeneration through oxidative stress, proteostasis, and immune responses ^49^. Other convergent DEGs like Neuregulin 1 (NRG1), Low-Density Lipoprotein Receptor Class A Domain-Containing 4 (LDLRAD4), Spectrin Repeat Containing Nuclear Envelope Protein 1 (SYNE1), Kin of IRRE-Like Protein 1 (KIRREL1), Rho GTPase Activating Protein 22 (ARHGAP22), Nuclear Factor I X (NFIX), Ectodermal-Neural Cortex 1 (ENC1), SLIT-ROBO Rho GTPase Activating Protein 3 (SRGAP3), Zinc Finger Protein 365 (ZNF365), Mediator Complex Subunit 12 Like (MED12L), Neuronal Cell Adhesion Molecule (NRCAM), Tropomyosin 4 (TPM4), Sec1 Family Domain-Containing Protein 2 (SCFD2), HIVEP Zinc Finger 1 (HIVEP1), Aryl Hydrocarbon Receptor Nuclear Translocator 2 (ARNT2), C-Type Lectin Domain Containing 16A (CLEC16A), Fos-Related Antigen 2 (FOSL2), Calcium Voltage-Gated Channel Auxiliary Subunit Beta 4 (CACNB4) may influence neurodegeneration through neuronal development, synaptic function, cytoskeletal organization, immune signaling, and transcriptional regulation.

Next, DEGs shared between interneurons and our human 3D *in vitro* brain model demonstrate alterations in HSP90AB1 and INSR that are directly linked to Alzheimer’s disease (AD) ^50,51^. HSP90AB1, a molecular chaperone, plays a role in protein folding and tau stabilization, which can influence tau aggregation in AD pathology. INSR (insulin receptor) drives brain glucose metabolism, which is associated with cognitive decline and amyloid-beta accumulation in AD ^52,53^.

Microglia cells had convergent DEGs directly linked to Alzheimer’s disease, such as STIP1, DTL, and indirectly, CTSB ^54–56^. Other DEGs commonly associated with epigenetic regulation of histones or DNA methylation are UHRF1, ATAD5, NSD2, and SMC4 ^57–59^. Given that microglia are believed to be at the core of Alzheimer’s disease onset and progression, these findings suggest that injury-induced microglial mitochondrial dysfunction and transcriptional alterations may contribute to the early priming of neuroinflammatory and neurodegenerative pathways, potentially accelerating disease progression in vulnerable individuals.

At last, some DEGs shared between astrocytes and our human 3D *in vitro* brain model play roles in transcription regulation (RFX2, BCL6, MED12L, KNDC1), cytoskeletal organization (SYNE1, BCAR3), ubiquitination (LRSAM1, UBE2H), DNA repair (BRIP1, FANCA, GINS3), metabolic transport (SLC7A11, SLC25A18, PSAT1), and signaling (ABR, ARHGAP39, ARHGAP26, LTBP1) ^60–63^. Though not directly connected to Alzheimer’s disease (AD), some DEGs like CTSB (cathepsin B), SLC7A11 (glutathione metabolism), and BCL6 (inflammation)^49^ may facilitate neurodegeneration through oxidative stress, proteostasis, and immune responses.

While studying convergence between human patients and our human 3D *in vitro* brain tissue model, we have discovered two emerging patterns: cell-specific mitochondria dysfunction and a link to neurodegeneration.

### Fabrication and validation of fluorescently labeled mitochondria for cell-specific analysis in brain injury

To further dissect the cell-specific mitochondria role in injury-induced neurodegeneration, we inserted genetic information for the expression of three distinct fluorescent tags under COX8A mitochondrial complex IV protein promoter in human brain cell types – human-induced neuronal stem-derived neurons, primary astrocytes, and immortalized HMC3 microglia (Figure S4 A-D).

Addgene provided two of the three fluorescent plasmids (EGFP and dsRED2) under the COX8A promoter (Figure S4B). We used VectorBuilder to implement a BFP2 fluorescent protein into the published backbone of mitochondria-targeted fluorescent plasmid under the COX8A promoter (Figure S4B) ^64^. We selected COX8A for all cell types due to its continuous presence and abundance in the mitochondria.

We first performed fluorescent intensity vs. viability experiments through various multiplicities of infection (MOI) (Figure S4C). Cell lines with the best viability and signal were selected. MOI40 was selected for mtdsRED2-iNSCs, MOI40 was selected for mtEGFP-astrocytes, and MOI10 was selected from mtBFP2-microglia. After transduction and expansion, we selected cells positive with target fluorophore using the Fluorescent-Activated Cell Sorter (FACS) (Figure S4D) and finally expanded cell lines for storage and experimentation. Please see the detailed description of the transduction protocol in the methods section.

Next, to validate the efficiency and specificity of the transduction, we co-stained 2D monocultures of the brain cells with mitochondria-specific marker Mitochondrially Encoded Cytochrome C Oxidase I (MTCO1) to ensure the fluorescent tags were mitochondria-specific. (Figure 3A). The images from panel A demonstrate precise co-localization of the mitochondria fluorescent tag with MTCO1 staining. Next, we isolated mitochondria from 2D monoculture dishes (using an Invitrogen mitochondria isolation kit from tissues) and human 3D *in vitro* brain models to validate mitochondria-specific protein markers via Western Blot analysis. Figure 4B displays cell-specific western blot stains of Phosphorylated Dynamin 1 Like / Dynamin 1 Like (pDRP1/DRP1), Mitochondrially Encoded Cytochrome C Oxidase I (MTCO1), Voltage Dependent Anion Channel 1 (VDAC), and Translocase Of Outer Mitochondrial Membrane 20 (TOMM20). The results demonstrate high OXPHOS activity in neuronal monocultures and tricultures (MTCO1 expression) and high glycolytic activity and mitochondrial fission in astrocytes (VDAC1 expression and pDRP1/DRP1). Next, we stained tricultures with cell-specific cytoplasmic marker Tubulin Beta 3 Class III (Tuj1) for neurons, Aldehyde Dehydrogenase 1 Family Member A1 (ALDH1A1) for astrocytes, and Transmembrane Protein 119 (TMEM119) for microglia to ensure the correct fluorescent colors were expressed in the corresponding cell type (Figure 3C). The fluorescent images demonstrated that integrated fluorescent tags were expressed in specific cell types.

**Figure 3.**
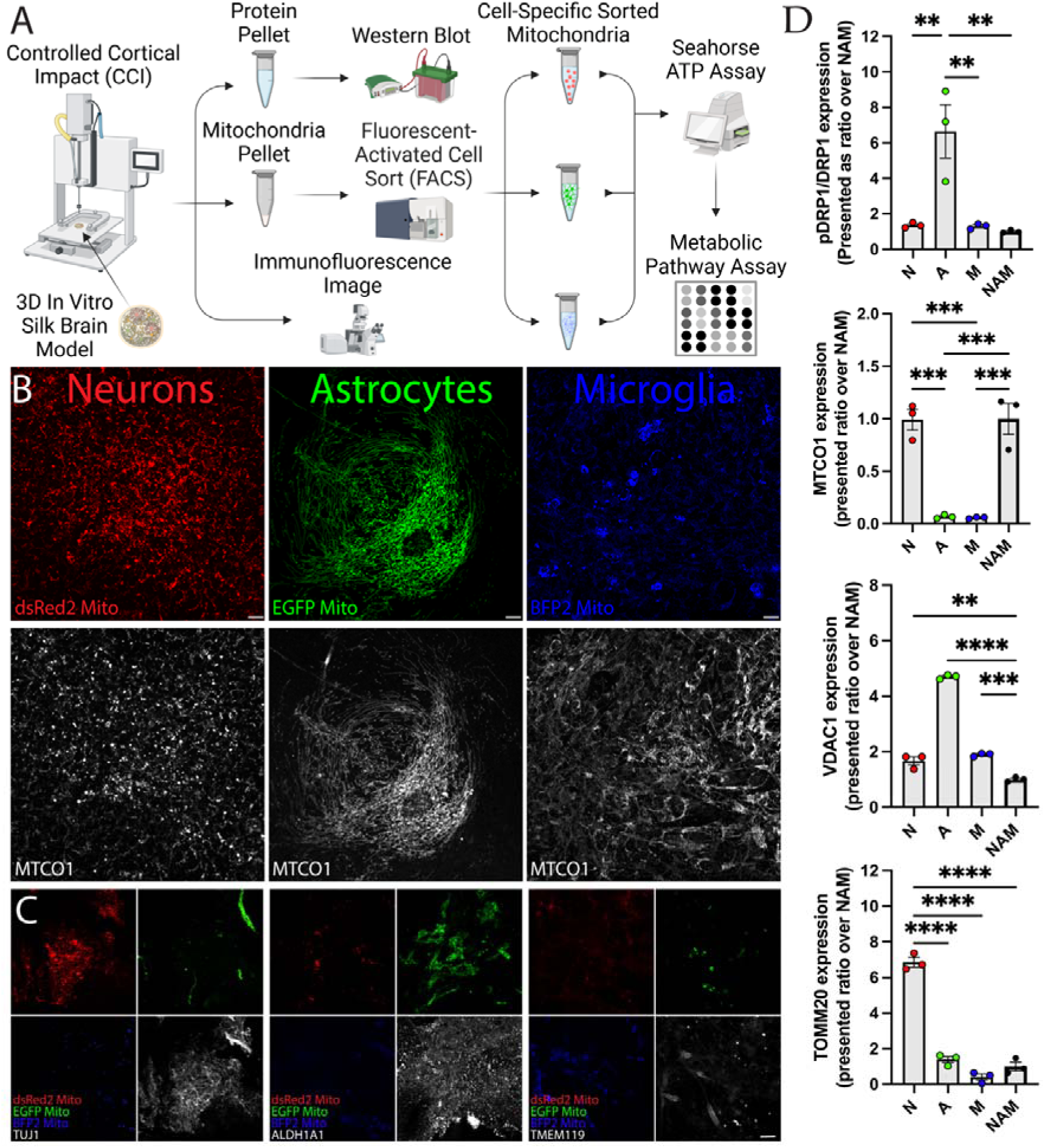
Molecular characterization and validation of the lentiviral transduction of the brain cells. (A) Schematic representation of the experimental designs. (B) Confocal images of 2D monocultures of mtdsRED2-Neurons, mtEGFP-Astrocytes, and mtBFP2-Microglia counterstained with MTCO1 (mitochondrially encoded cytochrome C oxidase I). Scale bar: 10 μm. (C) Confocal images of novel fluorescent cell lines grown for 5 weeks in 3D silk scaffolds, stained with a neuron (Tuj1-b3 tubulin), astrocyte (ALDH1A1 - Aldehyde Dehydrogenase 1 Family Member A1), and microglia-specific markers (TMEM119 – transmembrane protein 119) to validate cell-specific staining. Scale bar: 20 μm. (D) Mitochondria-specific markers analysis using Western blots of mitochondria isolated from 2D monocultures and NAM tricultures. pDRP1/DRP1 (Dynamin-related protein 1 phosphorylated at Ser 616 over total DRP1 protein) as a marker of mitochondria fission; MTCO1 as a marker of OXPHOS; VDAC1 (voltage-dependent anion channel 1) as a marker of glycolysis, and TOMM20 (translocase of outer mitochondria membrane 20) as a generic mitochondria marker. Data presented as mean±SEM of at least three independent experiments with n=3 for sham groups and n=4 scaffolds for injury groups in each independent experiment. * indicates a significant difference with p<0.05. One-way ANOVA with Tukey’s post-hoc test was used to determine the difference between control and experimental groups. The ROUT outlier analysis method was used to exclude statistical outliers. Data were normalized to sham at each time point.

**Figure 4.**
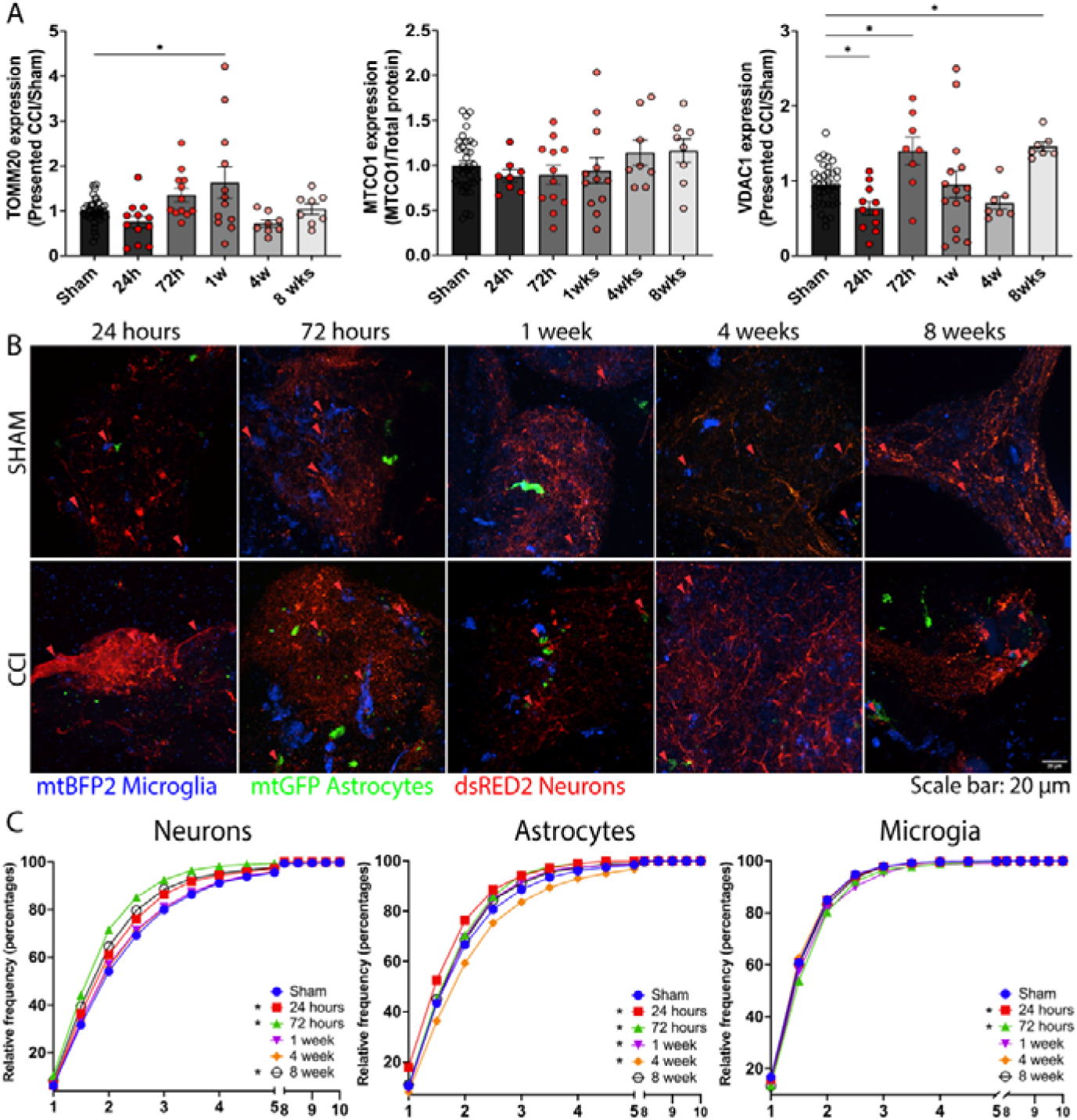
Acute and Chronic Cell-specific mitochondria fragmentation in response to moderate injury. A human 3D *in vitro* model composed of mtDsRED2-Neurons, mtEGFP-Astrocytes, and mtBFP2-HMC3 microglia matured in 3D for 6 weeks prior to being exposed to controlled cortical impact injury of moderate severity (6 m/s speed, 0.6 mm penetration, 3 mm tip) assessed 24 hours, 72 hours, 1 week, 4 weeks and 8 weeks post-injury. (A) Mitochondria-specific markers were assessed using Western blots: TOMM20 (translocase of outer mitochondria membrane 20) was used as a generic mitochondria marker; MTCO1 as a marker of OXPHOS; VDAC1 (voltage-dependent anion channel 1) as a marker of glycolysis. Data presented as mean±SEM of at least three independent experiments with n=3 for sham groups and n=4 scaffolds for injury groups. * indicates a significant difference with p<0.05. One-way ANOVA with Tukey’s post-hoc test was used to determine the difference between control and experimental groups. The ROUT outlier analysis method was used to exclude statistical outliers. Data were normalized to sham at each time point. (B) Representative confocal images of mitochondria fluorescent 3D in vitro cultures. Scale bar: 20 μm. (C) Frequency distribution (%) of cell-specific mitochondria fluorescent signal over time. Data for this plot were collected from at least three independent experiments with n=3 for sham groups and n=4 scaffolds for injury groups in each independent experiment. * indicates a significant difference with p<0.05. One-way ANOVA with Tukey’s post-hoc test was used to determine the difference between control and experimental groups. Data were normalized to sham at each time point.

### Cell-specific mitochondria fragmentation reveals neuronal vulnerability post-injury

Next, using our novel human 3D *in vitro* brain model with fluorescently labeled mitochondria, we assessed mitochondrial health and fission dynamics over time (at 5-time points - 24h, 72h, 1wk, 4wk, and 8wk) following controlled cortical impact (CCI) injury of moderate intensity (6 m/s with 3 mm tip and 0.6 mm penetration) (Figure 4A). Western blot analysis of bulk mitochondrial-specific markers at acute and chronic post-injury time points revealed that TOMM20 expression remained largely unchanged, except for a transient increase at 1-week post-injury. Similarly, MTCO1 expression remained stable throughout the study. However, VDAC1 exhibited a marked glycolytic depression at 24 hours, followed by a significant upregulation at 72 hours, aligning with previous reports of early metabolic suppression and compensatory glycolytic activation in TBI ^65–67^. VDAC1 levels then remained stable until a significant increase at 8 weeks, suggesting a late-stage metabolic shift.

To further assess mitochondrial structural dynamics, we evaluated mitochondrial fragmentation across neuronal, astrocytic, and microglial populations in our 3D in vitro model using confocal imaging of dsRED2-labeled neurons, EGFP-labeled astrocytes, and BFP2-labeled microglia (Figure 4B). Mitochondrial aspect ratio (length over width ratio) analysis (Figure 4C) revealed that all cell types exhibited substantial mitochondrial fragmentation at 24 hours post-injury. However, the recovery patterns varied across cell types. Neuronal mitochondria remained highly fragmented for up to 8 weeks, with only partial recovery observed between 72 hours and 8 weeks. Astrocytic mitochondria exhibited progressive fragmentation up to 4 weeks post-injury, followed by complete recovery at 8 weeks, indicating a delayed but reversible structural response. In contrast, microglial mitochondria demonstrated only transient fragmentation at 24 and 72 hours, with full recovery by 1 week, suggesting a more rapid adaptation to injury. These findings highlight neuronal and astrocytic mitochondria as the most vulnerable to persistent structural deficits.

### Cell-specific mitochondrial bioenergetics reveal divergent responses to injury

Mitochondrial fragmentation analysis provides critical insights into structural changes associated with bioenergetic function; however, it does not directly assess whether cells generate sufficient ATP to meet metabolic demands. To address this, we isolated mitochondria from scaffolds at five-time points—24h, 72h, 1w, 4w, and 8w—followed by Fluorescence-Activated Cell Sorting (FACS) to separate mitochondria based on cell type-specific fluorescence. Freshly sorted mitochondria were then assessed for bioenergetic function using the Seahorse ATP Rate Assay to measure oxygen consumption rate (OCR) and extracellular acidification rate (ECAR), followed by a Ray Biotech Metabolite Assay to analyze metabolic alterations (Figure 5A-B).

**Figure 5.**
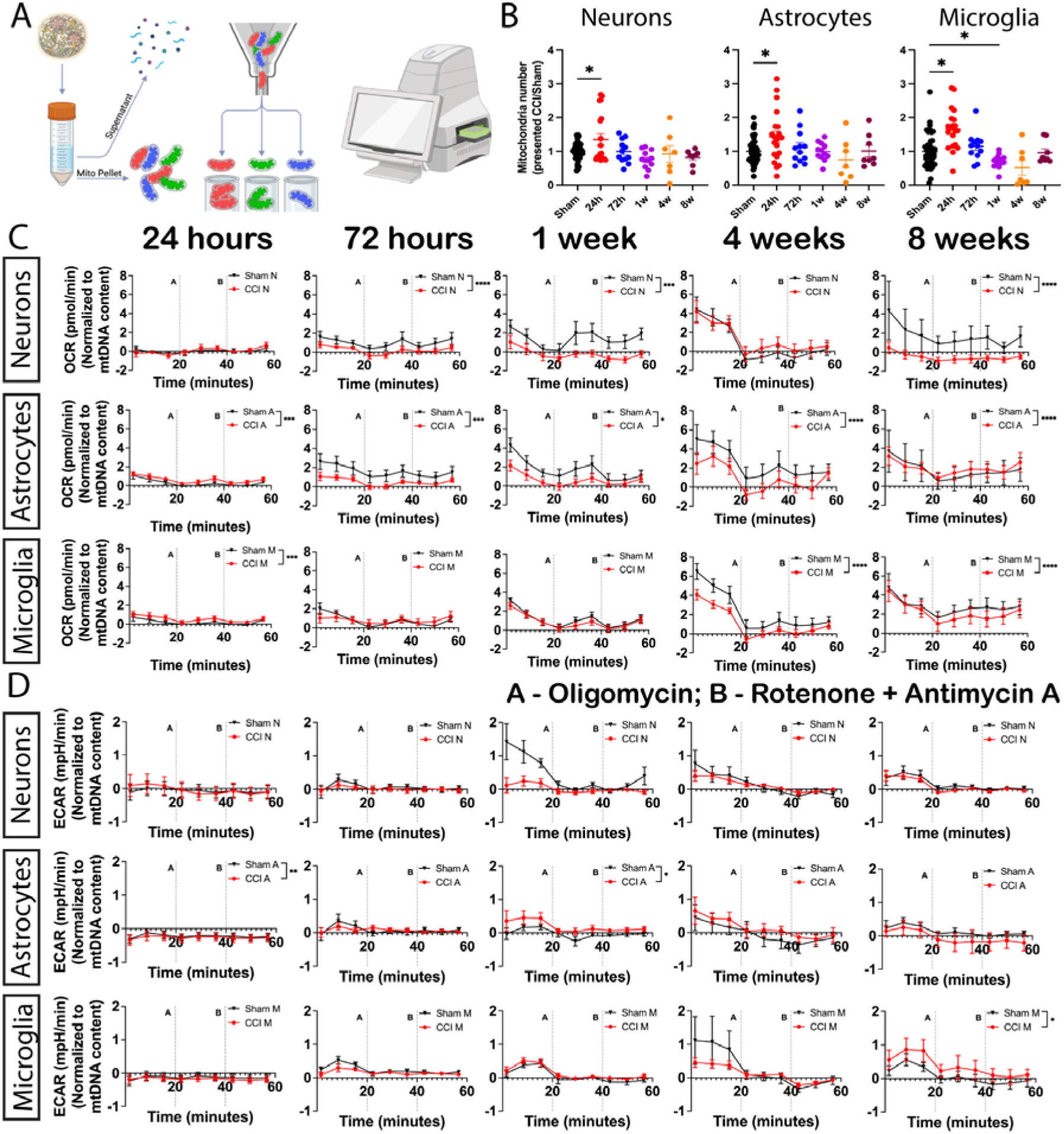
Cell-specific mitochondrial bioenergetics reveal divergent responses to injury. (A) Schematic representation of mitochondria sorting experiments followed by measurement of mitochondria bioenergetic function (B) The total number of cell-specific mitochondria sorted at each time point after injury. Data presented as mean±SEM of at least three independent experiments with n=3 for sham groups and n=4 scaffolds for injury groups in each independent experiment. * indicates a significant difference with p<0.05. One-way ANOVA with Tukey’s post-hoc test determined the difference between control and experimental groups. The ROUT outlier analysis method was used to exclude statistical outliers. Data were normalized to sham at each time point. (C) Oxygen Consumption Rate (OCR) and (D) Extracellular acidification rates (ECAR) were measured with real-time ATP rate assays on freshly sorted mitochondria 24, 72 hours, 1, 4, and 8 weeks post-injury. Data presented as mean±SEM of at least three independent experiments with n=3 for sham groups and n=4 scaffolds for injury groups. *, **, *** indicates a significant difference with p<0.05, p<0.01, p<0.001 respectively. Repetitive measures ANOVA with Tukey’s post-hoc test was used to determine the difference between control and experimental groups. The ROUT outlier analysis method was used to exclude statistical outliers. Presented OCR and ECAR rates were normalized to mitochondria DNA content.

Neuronal mitochondria exhibited a significant drop in OCR by 72 hours, followed by partial recovery at 4 weeks before declining again at 8 weeks, indicating prolonged bioenergetic instability (Figure 5B). Astrocytic mitochondria showed an initial increase in OCR acutely post-injury, followed by a steady decline until 4 weeks, after which levels stabilized. In contrast, microglial mitochondria demonstrated a slight acute increase in OCR, remained stable until 1 week, and then exhibited a significant reduction by 8 weeks. ECAR measurements at 24 and 72 hours were below detection limits across all cell types (Figure 5D). Astrocytic mitochondria displayed a transient increase in ECAR at 1 week, which significantly declined at 8 weeks, whereas microglial mitochondria maintained stable proton pumping capacity until a marked increase at 8 weeks. These findings reveal cell-specific metabolic vulnerabilities, with neuronal mitochondria experiencing persistent energy deficits, astrocytes exhibiting delayed but progressive dysfunction, and microglia undergoing late-stage metabolic reprogramming. At 20 minutes, we added Oligomycin; we added Rotenone + Antimycin A at 40 minutes. P value: * = .05| ** = .01| *** = .001|

### Divergent mitochondria metabolic profiles in brain cells following injury

Despite decades-long assumptions that mitochondria primarily function as cellular powerhouses, emerging evidence highlights their equally critical role in metabolic regulation, transcriptional control, and intracellular signaling, all of which are essential in the cellular response to injury and the progression of neurodegeneration. Thus, we next evaluated cell-specific metabolic alterations.

6A–C illustrate cell-type-specific metabolic alterations in neuronal, astrocytic, and microglial mitochondria at 24 hours and 8 weeks post-injury, as assessed by the Ray Biotech Metabolite Assay, with metabolite interaction networks analyzed via StringDB functional enrichment. These findings reveal distinct mitochondrial metabolic adaptations in response to injury, highlighting divergent mechanisms of recovery, dysfunction, and long-term metabolic reprogramming across brain cell types.

Neuronal mitochondria exhibited acute metabolic dysfunction at 24 hours post-injury, characterized by substantial alterations in pathways linked to cellular recovery, including nicotinate, nicotinamide, pyrimidine, purine, and riboflavin metabolism. By 8 weeks, these pathways return to baseline, suggesting metabolic resilience and recovery. However, early disruptions in sphingolipid metabolism and glycans degradation, key processes in post-translational protein modification, suggest transient impairments in protein homeostasis. Critical regulatory hubs identified in Figure 6C, including SIRT2, SIRT3, NAMPT, and ARSA, further emphasize mitochondrial involvement in coordinating metabolic recovery.

**Figure 6.**
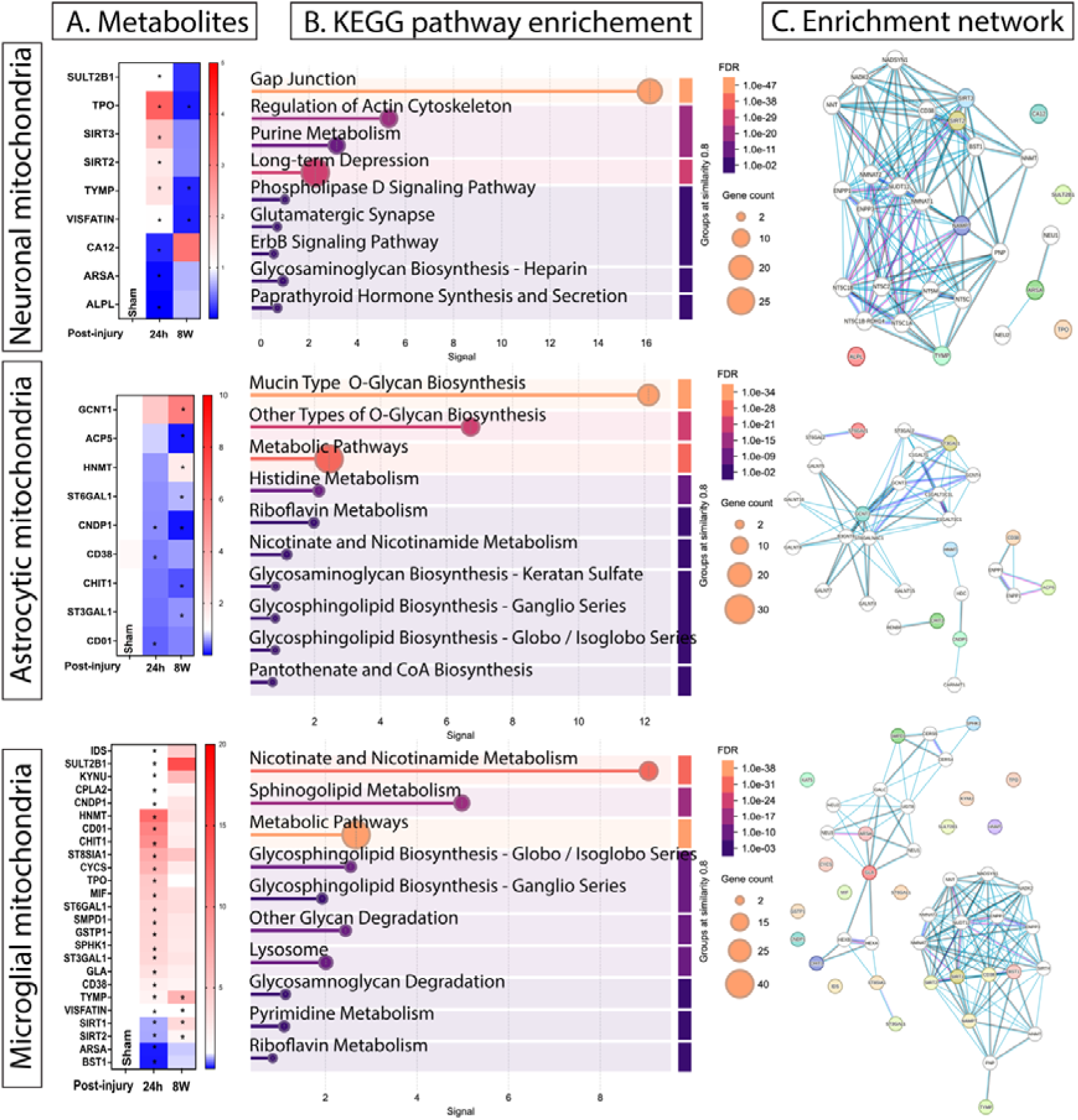
Divergent mitochondria metabolic profiles in brain cells following injury. Mitochondria that was stored after the bioenergetic assessment was used to determine metabolic alterations at acute (24 hours) and chronic (8 weeks) time points using the RayBiotech Metabolic panel. (A) Differentially produced metabolites Mitochondria from 3 independent experiments were assessed, with n=3 for sham groups and n=4 scaffolds for injury groups in each independent experiment. Data presented as mean±SEM of at least three independent experiments. * indicates a significant difference with p<0.05. Two-way ANOVA with Tukey’s post-hoc test determined the difference between the control and experimental groups. The ROUT outlier analysis method was used to exclude statistical outliers. Data were normalized to sham at each time point. (B) KEGG Pathway enrichment analysis isolated in panel A metabolites that were differentially produced. (C) Enrichment network corresponding to cell-specific metabolites from Panel A.

Astrocytic mitochondria showed no significant metabolic alterations at 24 hours, but by 8 weeks, they exhibited a profound decline in metabolic activity, particularly in pathways essential for O-glycans, glycosaminoglycans, glycophospholipids, and glycosphingolipid biosynthesis. This mirrors the progressive impairment in astrocyte-mediated metabolic support, potentially affecting neuronal function and long-term homeostasis. Like neuronal mitochondria, astrocytic mitochondria exhibit decreased pantothenate and CoA biosynthesis, which is crucial for hexosamine biosynthetic pathways regulating protein glycosylation and sialylation. However, unlike neuronal mitochondria, the astrocytic mitochondrial enrichment network is fragmented, reflecting a loss of metabolic coordination that may contribute to sustained dysfunction.

Microglial mitochondria, in contrast, show a rapid metabolic upregulation at 24 hours, with several metabolites remaining elevated at 8 weeks, including TYMP, VISFATIN, SIRT1, and SIRT2. These findings indicate that microglial mitochondria sustain an altered metabolic state, likely linked to prolonged inflammatory activation. Similar to neuronal and astrocytic mitochondria, microglial mitochondria exhibit alterations in sphingolipid, glycosphingolipid, and glycophospholipid metabolism, which are critical for membrane homeostasis and post-translational modifications. The enrichment network reveals two distinct microglial mitochondrial clusters—one highly interconnected network centered around SIRT1, SIRT2, CD38, and BST1, and a second, more fragmented network with multiple disconnected metabolic markers. This suggests that microglial mitochondrial metabolism remains in a prolonged state of dysregulation, likely contributing to chronic neuroinflammation and neurodegenerative processes following injury.

### Temporal Progression of Alzheimer’s-Like Neurodegenerative Markers Following Injury

The final question we investigated was whether a temporal link exists between injury-induced mitochondrial bioenergetic and metabolic dysfunction and the progression of AD-like pathology. To address this, we assessed neuronal network integrity (Figure 7A) alongside well-established neurodegenerative markers associated with Alzheimer’s disease (Figure 7B). The results indicate that key hallmarks of Alzheimer’s pathology, including Aβ42/40 ratio, pTau (AT8), TDP-43, APP, connexin 43, annexin A2, Neurogranin, and NCAM-1, exhibit injury-induced alterations at acute time points, with all intracellular markers beginning to increase by 72 hours post-injury (coincided with neuronal OCR bioenergetic decay).

**Figure 7.**
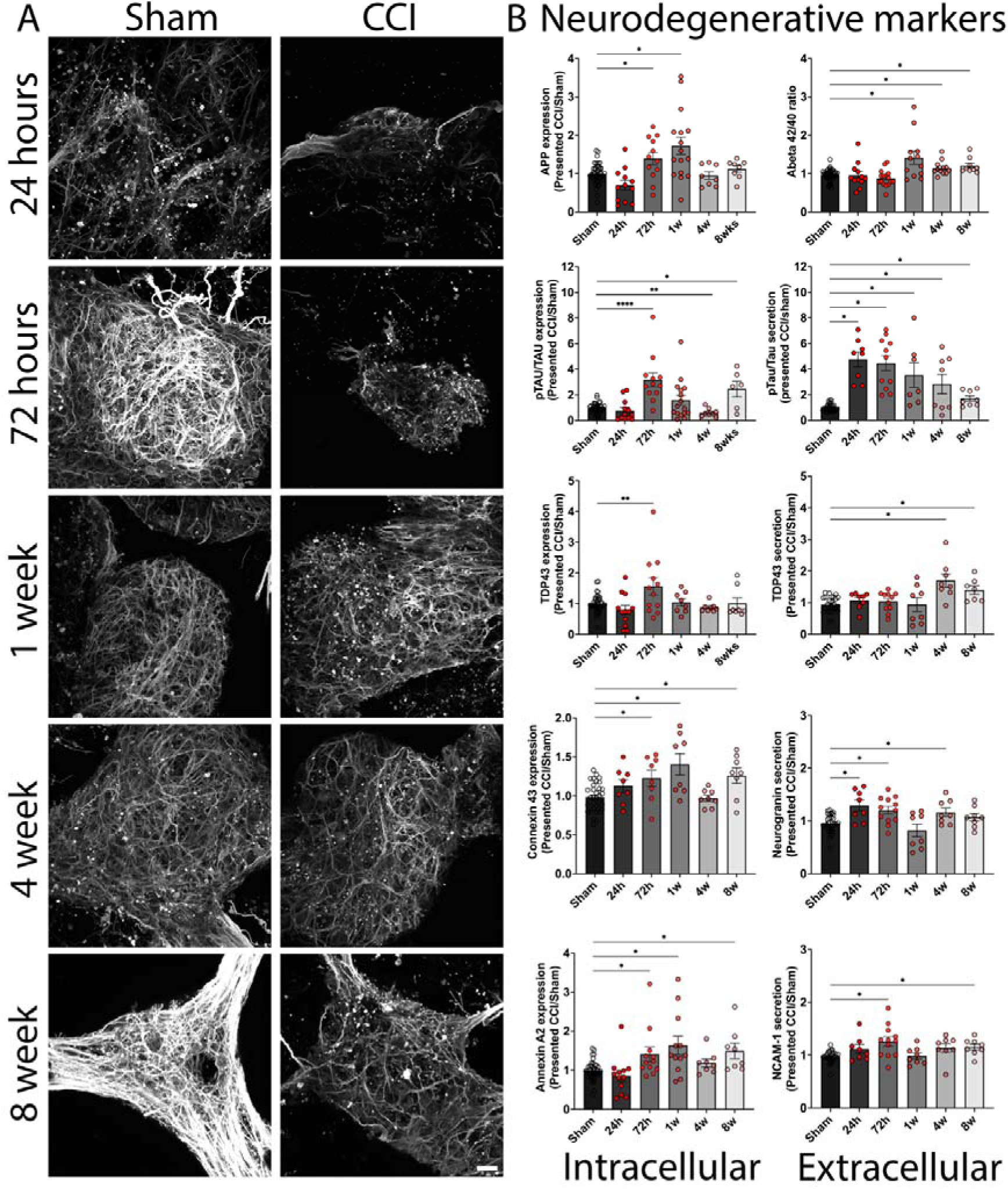
Mitochondria dysfunction in association with injury-induced AD-like neurodegenerative phenotype. (A) Displays representative confocal images of the 3D triculture scaffold of Tuj1 neuronal membrane stain over time for the sham and CCI groups. Scale bar: 10 μm. (B) Intracellular neurodegenerative markers were assessed by Western blot, while extracellular markers were assessed with a 9-plex neurodegeneration panel at acute (24 and 72 hours) and chronic (1, 4, and 8 weeks) post-injury time points. Intracellular markers: Phosphorylated Tau (AT8 at Ser202 and T205) presented over total Tau; TAR DNA Binding Protein (TDP43), amyloid precursor protein (APP), Gap Junction Protein Alpha 1 (Connexin43), Annexin A2 (ANXA2); Extracellular markers: Amyloid Beta 42 presented over Amyloid-Beta 40; phosphorylated Tau (AT8 at Ser202 and T205) over Tau, TDP43, Neurogranin, NCAM-1 (with FGF-21 and KLK6 not detected). Data presented as mean±SEM of at least three independent experiments with n=3 for sham groups and n=4 scaffolds for injury groups in each independent experiment. \**, **, \*\*\** indicates a significant difference with p<0.05, p<0.01, p<0.001 respectively. One-way ANOVA with Tukey’s post-hoc test determined the difference between control and experimental groups. The ROUT outlier analysis method was used to exclude statistical outliers. Data were normalized to sham at each time point.

Intracellular APP, a key precursor of Aβ, exhibited a significant increase from 72 hours to 1 week post-injury before returning to baseline levels by 4 weeks. In parallel, the Aβ42/40 ratio—a hallmark indicator of amyloidogenic processing in Alzheimer’s disease— showed a marked elevation at 1 week, which remained persistently upregulated at 4 and 8 weeks, indicating a sustained shift toward amyloidogenic progression following injury at chronic time points.

Intracellular Phospho-Tau (AT8) shows an initial increase at 72 hours, returns to baseline at 4 weeks, and becomes upregulated again at 8 weeks (follows the neuronal OCR pattern), while extracellular showed consistent upregulation up to 8 weeks, indicating the presence of a chronic neurodegenerative process despite apparent neuronal network recovery in Figure 7A. Intracellular TDP-43 follows a similar pattern, with upregulation observed at 72 hours post-injury, while extracellular was detected only at chronic time points. Intracellular Connexin 43 ^68,69^ and Annexin A2^70–72^, both of which are linked to Alzheimer’s disease, exhibit a similar temporal pattern, with upregulation at 72 hours, 1 week, and 8 weeks, followed by a transient decline at 4 weeks. Finally, extracellular neurogranin demonstrated acute increase (24 and 72 hours), with a brief spike at 4 weeks, and NCAM-1 first upregulated at 72 hours and 4 weeks, showing inconsistent transient changes. These findings establish a clear timeline in which mitochondrial dysfunction precedes and potentially drives progressive neurodegenerative changes, reinforcing the role of energy failure as a key contributor to injury-induced AD-like pathology.

## Discussion

Epidemiological studies demonstrated a strong link between traumatic brain injury (TBI) and an increased risk of neurodegenerative disorders, particularly Alzheimer’s disease (AD) ^5–8^, with elevated risk more than fourfold ^8^. Postmortem analyses of TBI patients reveal hallmark AD pathologies, including amyloid-beta plaques and hyperphosphorylated Tau, suggesting that even a single injury may predispose individuals to neurodegeneration ^8,9^. However, the mechanisms underlying this relationship remain poorly understood.

Mitochondrial dysfunction has emerged as a key early driver of injury-induced neurodegeneration ^6,10,73–75^, disrupting ATP production ^27,28,76,77^, calcium homeostasis, and oxidative balance ^11,78^ - conditions that not only impair neuronal function but may also promote pathological protein aggregation ^77,79,80^ and chronic inflammation ^81–84^, two central features of AD progression. Additionally, mitochondrial impairment affects lipid metabolism and glycosylation, both of which play crucial roles in cellular recovery and have been implicated in AD pathology ^85,86^. While mitochondria exhibit distinct phenotypic profiles across different brain cell types, it remains unknown whether these differences influence their response to trauma or contribute to long-term neurodegeneration ^87–93^. Addressing this gap is critical to understanding how TBI sets the stage for AD and identifying potential therapeutic targets to mitigate post-injury neurodegeneration.

Previous 24-hour transcriptomic analysis ^30^ showed extra and intracellular mitochondria dysfunction; here, we further corroborated these results with extended bulk transcriptomic analysis at 6- and 48-hours post-injury, indicating drastic changes to OXPHOS and mitochondria maintenance pathways (Figure 1). While previous studies, including our own, have established overall mitochondrial dysfunction, none have delineated cell-specific mitochondrial alterations. One study suggested acute, brain cell-specific mitochondrial dysfunction post-TBI using non-invasive metabolic imaging and secreted miRNA ^94^; however, no prior human cell or patient-based brain injury study ^16,17,95^ has comprehensively characterized the cell-specific bioenergetic and metabolic dysfunction of mitochondria across both acute and chronic time points or directly linked these alterations to injury-induced neurodegeneration.

Following brain injury, mitochondrial gene expression undergoes a dynamic shift, initially engaging compensatory mechanisms to sustain ATP production and mitigate oxidative stress (Figure 1-2). While early transcriptomic changes suggest a temporary recovery, long-term bioenergetic assessments reveal a sustained decline in mitochondrial function across all cell types. This persistent dysfunction challenges the notion of metabolic homeostasis, indicating that mitochondrial impairment is not a transient response but a prolonged pathological feature that may actively drive injury-induced neurodegeneration (Figures 1, 2, and 5) ^96–98^.

Our further cell-specific transcriptomic analysis between human TBI patients and our 3D in vitro brain injury model reveals a striking convergence in cell-specific mitochondrial dysfunction (Figure 2). This supports the hypothesis that metabolic disturbances are central to injury-induced neurodegeneration. The data show that while neurons primarily attempt to restore synaptic function post-injury, they also exhibit dysregulation of MAPK signaling, a key regulator of mitochondrial activity, oxidative stress, and apoptosis. In contrast, astrocytes and microglia demonstrate profound metabolic shifts, with early disruptions in OXPHOS and ATP production, followed by inflammatory activation and alterations in lipid metabolism. In addition to mitochondrial dysfunction markers, we identified multiple genes linked to Alzheimer’s disease (AD) across different brain cell types, reinforcing the potential role of mitochondrial impairment in driving injury-related neurodegeneration. Microglia exhibited alterations in STIP1, DTL, and CTSB, genes associated with AD ^54–56^, alongside changes in epigenetic regulators ^57–59^, Potentially influencing long-term neuroinflammatory responses, astrocytes displayed metabolic and transcriptional dysregulation, while excitatory neurons showed disruptions in CAMK2D and MAOB, which affect synaptic plasticity and oxidative stress, both hallmarks of AD ^44,45,47,99,100^. The convergence of mitochondrial dysfunction, metabolic stress, and neurodegenerative signatures across human patients and our in vitro model underscores the importance of mitochondrial health in determining long-term injury outcomes. These findings suggest that mitochondrial impairment may serve as an early driver of injury-induced neurodegeneration, linking TBI to the onset of AD-like pathology. Additionally, this analysis reinforces the physiological relevance of our in vitro model.

Using the initial analysis comparing human data with our bulk transcriptomics, we hypothesized that neuronal mitochondria experience severe damage and require immediate support from glial cells to restore function. To investigate this, we developed a unique brain injury model that enables cell-specific live mitochondrial research (Figure 3-7).

An integrated analysis of cell-specific transcriptomics, mitochondrial fragmentation, bioenergetics, and metabolic fluctuations demonstrated a clear mechanistic link between injury-induced mitochondrial dysfunction and the emergence of neurodegenerative markers. Confocal imaging of mitochondrial health revealed that while outer membrane integrity (TOMM20) remained stable, VDAC1 expression exhibited an acute drop at 24 hours, aligning with known glycolytic suppression in TBI patients ^65–67,101^. Mitochondrial bioenergetic assessments further supported this, as neuronal OCR dropped substantially by 72 hours, marking the first significant decline in mitochondrial respiration. This decline coincided with a sharp increase in neurodegenerative markers, including Aβ 42/40, pTau, APP, and TDP-43, suggesting that mitochondrial dysfunction may directly contribute to early neurodegenerative processes. The sustained neuronal mitochondrial fragmentation for up to 8 weeks, along with fluctuations in OCR recovery and subsequent decline, indicates that neurons fail to fully restore metabolic homeostasis, making them highly susceptible to long-term degeneration.

Astrocytic and microglial mitochondria displayed distinct metabolic trajectories, with astrocytes showing minimal acute metabolic disruptions but developing chronic impairments in glycan biosynthesis and lipid metabolism by 8 weeks, as seen in our metabolic panels (Figure 6). Conversely, microglial mitochondria initially upregulated metabolic pathways, including markers such as SIRT1 and SIRT2, but exhibited persistent inflammation-associated metabolic activity at chronic time points. Notably, at 72 hours, microglial OCR remained stable while astrocytes showed an increasing trend, contrasting with neuronal OCR decline, suggesting that while glial cells actively respond to metabolic demands, neurons experience an early energy crisis that coincides with neurodegenerative marker upregulation.

The Aβ42/40 ratio, a key hallmark of AD pathology, ^100,102,103^ Demonstrated a significant increase at chronic time points, mirroring mitochondrial metabolic instability. This pattern is highly relevant to Alzheimer’s disease (AD) pathology, where disruptions in Aβ metabolism contribute to synaptic toxicity and neuronal dysfunction, eventually leading to excessive Aβ accumulation and aggregation. This shift, combined with sustained pTau upregulation and persistent neuronal OCR instability, suggests that mitochondrial dysfunction after injury may create a permissive environment for AD-like neurodegeneration, potentially accelerating disease onset in predisposed individuals. These findings reinforce the growing body of evidence that TBI serves as a trigger for AD pathology, with mitochondrial failure playing a central role in the transition from acute injury to chronic neurodegeneration.

In summary, our study integrated cell-specific mitochondrial profiling with patient-derived transcriptomic data, providing unprecedented insight into how mitochondrial dysfunction progresses following TBI and the first comprehensive evidence that mitochondrial dysfunction is not a uniform event but instead manifests in a cell-type-specific manner, with neuronal mitochondria exhibiting the most prolonged metabolic instability and direct correlation with neurodegenerative marker upregulation. Our cell-specific mitochondrial analyses aligned with transcriptomic findings, reinforcing that neurons struggle with energy production, astrocytes fail to maintain metabolic support long-term, and microglia sustain chronic inflammatory activity. This confirms that neuronal bioenergetic failure is an early driver of neurodegeneration, while astrocytes and microglia contribute to disease progression over time. Our approach proved helpful in identifying mitochondrial dysfunction as a therapeutic target, emphasizing the need for interventions that stabilize neuronal metabolism early after injury to prevent long-term neurodegeneration.

### Limitations to the study

While our study provides unprecedented insights into cell-specific mitochondrial dysfunction and its role in injury-induced neurodegeneration, several limitations should be considered. First, our model lacks a vascular component and neuronal myelination, limiting our ability to assess how blood-brain barrier integrity and white matter injury contribute to metabolic dysfunction. The absence of these key elements may underestimate the impact of vascular and myelination deficits on long-term neurodegeneration. Second, our study utilized a single genetic background, restricting our ability to account for individual variability in mitochondrial resilience, immune responses, and neurodegenerative susceptibility. Future studies incorporating multiple donor-derived cell lines will be necessary to determine whether genetic differences influence mitochondrial responses to injury. Lastly, our study is limited to an 8-week post-injury period, providing insight into early and intermediate stages of injury progression but not capturing long-term neurodegenerative changes beyond this window. Extending the study duration would allow us to determine whether mitochondrial dysfunction persists or worsens over time, particularly in relation to AD-like pathology. Addressing these limitations in future research will enhance our understanding of how mitochondrial dysfunction contributes to chronic neurodegeneration following TBI.

## Authors contribution

VL conceived the project. SBSK and VL designed and interpreted the experiments and wrote the manuscript. SBSK and SPS performed scaffold seeding and sample maintenance (media changes). SBSK performed viral transductions, all in vitro experiments, injuries, and Raybiotech assays. SBSK and VL performed mitochondria isolation and FACS sorting. SPS and SBSK performed mitochondria bioenergetic function assessment. SPS, VL, and SBSK performed confocal imaging. SBSK, SPS, CA, and MT conducted Western Blots. VL performed ELISA. SBSK performed a multiplex panel. CA performed an aspect ratio analysis. SBSK and SPS prepared silk materials. SBSK performed RNA sequencing analysis.

## Acknowledgments

The authors thank the University of Cincinnati Start-Up Funds and Research Scholar Award, NIH (1R21AG085052-01A1), Cincinnati Children’s Hospital Bioimaging and Analysis Facility, and Research Flow Cytometry Facility.

## Methods

### Fabrication of 3D Silk Scaffolds

3D silk scaffolds were fabricated following established protocols ^30,104,105^. Bombyx mori cocoons were boiled in 0.02 M sodium carbonate for 30 minutes and then dried overnight under a fume hood. The dried silk fibers were dissolved in 9.3 M lithium bromide at 60 °C for 4 hours. To remove lithium bromide, the silk solution underwent dialysis in deionized water at room temperature for 72 hours using 3,500 molecular weight cut-off dialysis tubing (Invitrogen), with water changes every 12 hours. The dialyzed silk solution was centrifuged at 9,000 rpm for 20 minutes twice, and debris was removed using a 100 µm pore strainer. The concentration was adjusted to 6 mg/mL before casting into 10 cm dishes. To induce porosity, 400–500 µm sodium chloride (Sigma) particles were applied at a 1:2 (v/w) ratio. The solution was left to solidify at room temperature for 48 hours, followed by incubation at 60 °C for 1 hour to form silk sponges. Silk sponges underwent further dialysis in deionized water at room temperature for 48 hours, with six total water changes.

Scaffolds were cut into a 6 mm outer diameter and 2 mm inner diameter using biopsy punches (Integra) and trimmed to a height of 1.5 mm. Prior to cell seeding, scaffolds were autoclaved in deionized water for 20 minutes on a liquid cycle and equilibrated to room temperature. Scaffolds (n = 50 per condition) were placed in 6-well plates and incubated overnight at 37 °C in 7 mL of 10 µg/mL poly-L-ornithine (PLO) solution. Following three washes in distilled water at 5-minute intervals, scaffolds were incubated at 4 °C overnight in 0.5 mg/mL laminin (ThermoFisher 50-100-3381) prepared in phenol red-free Dulbecco’s Modified Eagle Medium/Nutrient Mixture (DMEM F-12, Gibco 21041025). Before use, scaffolds were equilibrated at 37 °C.

### Mycoplasma testing

All cell lines used in this project were either checked for mycoplasma before starting or were directly purchased from established vendors that provided certification for Mycoplasma testing.

### Mouse embryonic fibroblasts (MEFs)

Mouse embryonic fibroblasts (MEFs) were cultured as a feeder layer for human-induced neuronal stem cells (hiNSCs) following ATCC guidelines. MEFs at passages 1–4 were maintained in Dulbecco’s Modified Eagle Medium (DMEM) (1X) + GlutaMAX (ThermoFisher, 10569010) supplemented with 10% fetal bovine serum (FBS) (Gibco, 10-438-026) and 1% antibiotic-antimycotic (ThermoFisher, 15240062). Cells were incubated at 37 °C with 5% CO[, and media was changed every 3–4 days.

For seeding, 15 cm culture dishes were coated with 0.1% gelatin for 1 hour at room temperature before plating MEFs in 20 mL of MEF media. For expansion, cells were washed with phosphate-buffered saline (PBS), detached using 0.25% Trypsin-EDTA for 3 minutes at 37 °C, neutralized with MEF media, and centrifuged at 1,000 rpm for 5 minutes. Cells were split at a 1:3 ratio and passaged at 70–80% confluency.

To prepare MEFs as a feeder layer, cells were grown to 90–100% confluency and inactivated by incubation with 10 µg/mL Mitomycin C (Sigma, M4287-5X2MG) in MEF media at 37 °C for 3 hours. Following inactivation, cells were washed three times with PBS. If hiNSCs were seeded immediately, 20 mL of hiNSC media was added. If not used immediately, inactivated MEFs were maintained in 20 mL of MEF media with media changes every 3–4 days for up to 7 days.

### Human-induced neural stem cells (iNSCs) and mtRFP-iNSCs

Induced neural stem cells (iNSCs) were generated from dermis-derived human fibroblasts, originally derived from neonatal foreskin fibroblasts ^106^. Cells were maintained in KnockOut DMEM (ThermoFisher, 10829018) supplemented with 1% GlutaMAX (ThermoFisher, 35050061), 1% antibiotic-antimycotic (ThermoFisher, 15240062), 0.2% 2-Mercaptoethanol (ThermoFisher, 21985023), and 20% KnockOut Serum Replacement (KNSR) (ThermoFisher, 12-618-013). Cultures were incubated at 37 °C with 5% CO[, and media was refreshed every 1–2 days.

For seeding, complete iNSC media was prepared by supplementing 50 mL of iNSC media with 800 μL of 10 µg/mL basic fibroblast growth factor (bFGF) (ThermoFisher, PHG0261). Prior to seeding, MEF feeder plates were cultured until 90–100% confluency and inactivated with Mitomycin C. iNSCs were plated on inactivated MEF feeders in 20 mL of iNSC media.

For expansion, 15 cm culture dishes were coated with 2% Matrigel (Millipore Sigma CLS354234) diluted in DMEM F-12 (Gibco 21041025) for 1 hours. iNSCs were maintained until reaching 70–80% confluency. Cells were washed with PBS, detached using TrypLE at 37 °C for 1 minute, neutralized with iNSC media, and centrifuged at 3,000 rpm for 2 minutes. The cell pellet was resuspended by gentle pipetting (5 times) and split at a 1:20 ratio for further expansion. The same protocol was used for culturing fluorescently labeled iNSC lines.

### Human primary astrocytes and mtGFP-Astrocytes

Primary astrocytes were cultured in astrocyte growth media containing fetal bovine serum (FBS), astrocyte growth supplement, and Penicillin-Streptomycin (Pen/Strep) solution (ScienCell Research Laboratories, NC9707094). Cells were maintained at 37 °C with 5% CO[, and media was refreshed every 3–4 days.

For seeding, 15 cm culture dishes were coated with either 2.5 μL Poly-L-Lysine (Sigma, P4832) in 15 mL sterile distilled water for 1 hour (normal astrocytes) or 25 μL Poly-L-Lysine, 150 μL Laminin (ThermoFisher, 50-100-3381), and 15 mL sterile distilled water for 1 hour (mtGFP-labeled astrocytes). After coating, astrocytes were plated in 20 mL of astrocyte media.

For expansion, cells were washed with PBS, detached using 0.25% Trypsin-EDTA for 3 minutes at 37 °C, neutralized with astrocyte media, and centrifuged at 1,000 rpm for 5 minutes. Cells were resuspended and re-seeded at a density of 350,000 cells per 15 cm dish. Passaging was performed at 70–80% confluency.

### HMC3 microglia and mtBFP-Microglia

HMC3 human microglial cells were cultured in Eagle’s Minimum Essential Medium (EMEM) (ThermoFisher, 50-188-268FP) supplemented with 10% fetal bovine serum (FBS) (Gibco, 10-438-026) and 1% antibiotic-antimycotic (ThermoFisher, 15240062). Cultures were maintained at 37 °C with 5% CO[, and media was changed every 3–4 days.

For seeding, HMC3 cells were plated in 20 mL of HMC3 media. For expansion, cells were washed with PBS, detached using 0.25% Trypsin-EDTA for 3 minutes at 37 °C, neutralized with HMC3 media, and centrifuged at 1,000 rpm for 5 minutes. Cells were resuspended and re-seeded at a density of 350,000 cells per 15 cm dish. Passaging was performed at 70–80% confluency. The same protocol was used for subculturing fluorescently labeled cell lines.

### Lentivirus Transduction Protocol

#### Seeding Density

Seeding cells at different densities are required to quantify optimal confluency and cell growth of each cell type.

#### hiNSC

Coat plates with 250 microL of Matrigel diluted in 24 mL of DMEM F12. Reconstitute, add 2 mL per plate and wait 1 hour. Prepare hiNSC cells from cryovial or dishes. Count cells. Centrifuge cells for 2 min at 3000 rpm. Remove supernatant and add hiNSC media. Resuspend cell pellet 5 times. Add number of cells based on map and add sufficient media of each well to 2 mL. Wait 24 hours, image wells, and choose optimal seeding density.

#### Astrocyte

Coat plates with 2.5 microL of Poly-L-Lysine diluted in 12 mL of sterile UltraPure water. Add 2 mL per well and wait 1 hour. Prepare astrocyte cells from cryovial or dishes. Count cells. Centrifuge cells for 5 min at 1000 rpm. Remove supernatant and add astrocyte media. Resuspend cell pellet 5 times. Add number of cells based on map and add sufficient media of each well to 2 mL. Wait 24 hours, image wells, and choose optimal seeding density.

#### HMC3

Prepare HMC3 cells from cryovial or dishes. Count cells. Centrifuge cells for 5 min at 1000 rpm. Remove supernatant and add HMC3 media. Resuspend cell pellet 5 times. Add number of cells based on map and add sufficient media of each well to 2 mL. Wait 24 hours, image wells, and choose optimal seeding density.

#### Lentiviral Transduction

In order to quantify optimal transduction efficiency and cell growth of each cell type, transducing cells at different MOIs is required. Abided to BSL 2 Protocol link protocol

### Lentivirus Transduction

*hiNSC:* The hSYN-mito-dsRED2 plasmid (Addgene, #173069) was used to generate dsRED2-expressing iNSCs via lentiviral transduction.

*Day 1*: Corning 6-well plates were coated with Matrigel by adding 20 µL of Matrigel diluted in 2 mL of DMEM/F12 (Gibco 21041025) per well. After 1 hour at room temperature, the Matrigel solution was aspirated, and iNSCs were seeded at a density of 750,000 cells per well. Sufficient hiNSC media was added to achieve a final well volume of 2 mL. Plates were incubated overnight at 37 °C with 5% CO[. *Day 2*: Lentiviral cryovials were transferred from −80 °C to an ice bucket and thawed completely. hiNSC media was aspirated from each well, and lentiviral particles were added according to the viral titer and the desired multiplicity of infection (MOI). A fresh micropipette tip was used for each well. Media was adjusted to maintain a final volume of 2 mL per well. Control wells received 2 mL of hiNSC media without lentivirus. Plates were incubated overnight at 37 °C with 5% CO[. All equipment that came into contact with lentiviral particles was treated with bleach for 30 minutes before disposal. *Day 3*: After 24 hours, media was aspirated from each well using a new pipette tip for each well and disposed of in bleach. Wells were washed with 2 mL of PBS, and the wash solution was removed. Fresh iNSC media (2 mL) was added to each well, and plates were returned to the incubator at 37 °C with 5% CO[. Media changes were performed every 3–4 days.

*Astrocyte:* The pLV-mitoGFP plasmid (Plasmid #44385) was used to generate GFP-expressing astrocytes via lentiviral transduction.

*Day 1*: Corning 6-well plates were coated with Poly-L-Lysine (.333 microL of Poly-L-Lysine and 2 mL of UltraPure sterile water). After 1 hour at room temperature, Poly-L-Lysine solution was aspirated, and astrocytes were seeded at a density of 300,000 cells per well. Sufficient astrocyte media was added to achieve a final well volume of 2 mL. Plates were incubated overnight at 37 °C with 5% CO[. *Day 2*: The same procedure was conducted as Day 2 for iNSC transduction, but EGFP lentivirus and astrocyte media were used instead of DsRed2 lentivirus and iNSC media. *Day 3*: The same procedure was conducted as Day 2 for iNSC transduction, but used astrocyte media instead of iNSC media.

*HMC3:* The custom plasmid used VectorBuilder to implement a double COX8A promoter and EBFP2 into the plasmid backbone, published by Isamu Taiko and his group [14].

*Day 1*: Corning 6-well plates were not coated. HMC3 were seeded at a density of 300,000 cells per well. Sufficient astrocyte media was added to achieve a final well volume of 2 mL. Plates were incubated overnight at 37 °C with 5% CO[. *Day 2*: The same procedure was conducted as Day 2 for iNSC transduction, but EBFP2 lentivirus and HMC3 media were used instead of DsRed2 lentivirus and iNSC media. *Day 3*: The same procedure was conducted as Day 2 for iNSC transduction, but HMC3 media was used instead of iNSC media.

### 3D in vitro human triculture brain tissue model fabrication

3D *in vitro* human triculture brain tissue models were produced following the same protocol established in. RFP-expressing induced neural stem cells (RFP-INSCs), GFP-expressing astrocytes (GFP-Astros), and BFP-expressing human microglial clone 3 cells (BFP-HMC3) were cultured to near 100% confluency. Cells were lifted using monoculture methods and combined at a ratio of 2:0.5:0.1 million (RFP-INSCs:GFP-Astros:BFP-HMC3). The cell suspension was centrifuged at 1000 rpm for 5 minutes, and the supernatant was aspirated. Cells were resuspended in neurobasal medium (ThermoFisher 21-103-049) supplemented with 2% B-27 (ThermoFisher 17-504-044), 1% Anti-Anti (ThermoFisher 15240062), 1% GlutaMAX (ThermoFisher 35050061), and 1% astrocyte growth factors (ThermoFisher NC9746822).

Poly-L-ornithine (PLO)- and laminin-coated silk scaffolds were prepared in 96-well plates, and excess liquid was removed using a vacuum line. A 40 µL cell suspension (2:0.5:0.1 million cells) was seeded onto each 3D silk scaffold. Scaffolds were incubated at 37°C for 30 minutes to allow cell attachment, followed by the addition of 150 µL neurobasal medium per well. Scaffolds were incubated overnight at 37°C with 5% CO2.

The following day, cell-seeded scaffolds were transferred to new 96-well plates. A 100 µL collagen type I solution (ThermoFisher 344702001) was added to each scaffold, and the plates were incubated for 30 minutes at 37°C with 5% CO2. Subsequently, 150 µL neurobasal medium was added to each well, and the scaffolds were incubated overnight at 37°C with 5% CO2.

The next day, brain-like tissues were transferred to 48-well plates, each containing 1 mL neurobasal medium. The medium was changed every 3-4 days, and the plates were incubated at 37°C with 5% CO2. 3D in vitro silk brain models were ready 6 weeks post-seeding.

### Mitochondria Isolation

Mitochondria were isolated from 2D cell lines using the Mitochondria Isolation Kit for Cultured Cells (Thermo Fisher Scientific). Human microglial clone 3 cells (HMC3), astrocytes, and induced neural stem cells (hiNSC) were seeded independently in 6-well plates at densities of 350,000, 350,000, and 1,000,000 cells per well, respectively, and cultured for 3 days.

Cells were harvested, and mitochondria were isolated according to the manufacturer’s Option A protocol. Briefly, cells were centrifuged at 850 × g for 2 minutes. The supernatant was removed, and cells were resuspended in 800 µL Reagent A. The suspension was transferred to a 2.0 mL microcentrifuge tube, vortexed at medium speed for 5 seconds, and incubated on ice for exactly 2 minutes. 10 µL Reagent B was added, and the sample was vortexed at maximum speed for 5 seconds. After a 5-minute incubation on ice, with vortexing at maximum speed every minute, 800 µL Reagent C was added by gentle inversion. The tube was centrifuged at 700 × g for 10 minutes at 4°C with low braking. The supernatant was transferred to a 1.0 mL microcentrifuge tube and centrifuged at 12,000 × g for 15 minutes at 4°C. The supernatant was aspirated, and the mitochondrial pellet was resuspended in 100 µL RIPA buffer. Samples were stored at −80°C until Western blot analysis.

Use tweezers to transfer scaffolds to 2mL eppendorfs. Add 800µL of Mito Iso Reagent A. Squish and open the scaffold to remove cells. Vortex samples for 5 seconds on medium speed and incubate on ice for 2 minutes. Add 10µL of Mito Iso Reagent B and vortex on medium speed for 5 seconds. Incubate on ice. Vortex every minute for 5 minutes. Add 800µL of Mito Iso Reagent C to each sample. Invert 10 times (do not vortex). Remove scaffold. Centrifuge eppendorfs at 1000xg for 10 minutes at 4°C. Transfer supernatant to new 2mL eppendorfs. Throw out eppendorfs that only contain pellets (cell debris). Centrifuge supernatant Eppendorf at 12,000xg for 25 minutes at 4°C. Transfer supernatant to new 2mL Eppendorf (this is the protein fraction). Resuspend the pellet in 300µL of Seahorse Buffer (this is the mitochondria pellet).

### Seahorse ATP Rate Assay

The day before the assay, we prepared the Extracellular Flux Assay Pack by separating the XF Sensor Cartridge and the 96-well utility plate. Using a multichannel pipette, each well of the utility plate was filled with 200 µL of sterile water. After placing the pink protective guard on the plate, we inserted the sensor cartridge and incubated the assembly overnight at 37°C in a non-CO[ incubator to allow for sensor hydration.

On the day of the assay, we replaced the sterile water in the utility plate with 200 µL of Seahorse calibrant solution per well and returned the plate and cartridge to the non-CO[ incubator until use. For the assay media, we supplemented 50 mL of DMEM/F12 (Gibco 21041025) Phenol Red-free media with 0.5 mL each of pyruvate, glucose, and glutamine. We prepared the compound injections by diluting Oligomycin in 420 µL of Seahorse media and further diluting 300 µL of this solution into 2.7 mL of media. For Rotenone + Antimycin A, we used 540 µL of Seahorse media for the initial dilution, followed by the same 300 µL in a 2.7 mL step.

After isolating and sorting mitochondria by FACS, we resuspended the mitochondrial pellets in 180 µL of Seahorse assay media and transferred them into the assay plate. We then used Wave Desktop Software to designate sample groups, calibrate the cartridge, and run the assay for one hour. We normalized the resulting data to DNA content for each sample.

### DNA Quantification

After the Seahorse ATP Rate Assay, 20 µL of mitochondria sample was used for DNA quantification. 20 µL of Triton-X solution was added to each mitochondria sample, followed by incubation at room temperature (RT) for 10 minutes. Samples were then diluted 1:5 in TE buffer to optimize the fluorescence signal. A standard curve was generated using DNA lambda standards (2000–0 ng/mL). Samples and standards were mixed with 100µL of PicoGreen reagent in black 96-well plates and incubated in the dark for 5 minutes at RT. Fluorescence intensity was measured using a microplate reader (Excitation: 485nm, Emission: 538nm). DNA concentration is determined by interpolation from the standard curve and used to normalize mitochondrial bioenergetics data to DNA content (pg DNA/cell).

### Immunofluorescence

Transfer the scaffold to 48 well plates. Add 300µL of 4% PFA / 4% Sucrose and wait for 1 hour. Aspirate solution. Add 1mL of PBS and do 4 10-minute washes put on a shaker (RT). Aspirate PBS, Add 300µL of IF blocking solution, and incubate on a shaker (RT) for 1 hour. Aspirate IF blocking solution. Add 300µL of primary antibody (1:1000 dilution) and incubate on a rocker (4°C) overnight. Aspirate primary antibody. Add 1mL of PBS and put on a shaker (RT) for 10 minutes. Repeat 2 more times. Aspirate PBS. Cover the plate with aluminum foil (secondary is light-sensitive). Add 300µL of secondary antibody (1:500 dilution) and incubate on a shaker (RT) at slow speed for 1 hour. Aspirate secondary antibody. Add 1mL of PBS and put on a shaker (RT) for 10 minutes. Repeat 2 more times. Aspirate PBS. Add .5mL PBS. Store in 4°C or proceed with imaging.

*IF Primary*: MTCO1 Ms (Invitrogen REF 459600), Tuj1 (Anti-beta III Tubulin) Ms (Abcam ab78078), ALDH1A1 Rb (Invitrogen REF MA5-42586), TMEM119 Rb (Invitrogen REF PA5-119902). *IF Secondary*: Goat anti-Chicken IgY (H+L) (Invitrogen REF A32933), Goat anti-Mouse IgG (H+L) (Invitrogen REF A32728), Goat anti-Rabbit IgG (H+L) (Invitrogen A32733)

### RayBio Human Metabolic Pathways Antibody Array 1

Remove the kit from –20°C and let it thaw. Remove the Antibody Arrays from the plastic packaging. Place each membrane face-up into a 6-well incubation tray (ensure proper orientation). Add 2mL of RayBio Blocking Buffer to each well and incubate on a shaker (RT) at slow speed for 30 minutes. Aspirate the RayBio Blocking Buffer.

Pipette 1 mL of sample into each well and incubate on shaker (RT) at slow speed overnight. Aspirate sample from each well. Add 2mL of 1x Wash Buffer I to each well and incubate on shaker (RT) at slow speed for 5 minutes. Repeat 2 more times. Aspirate 1x Wash Buffer I. Add 2mL of 1x Wash Buffer II to each well and incubate on shaker (RT) at slow speed. Repeat 1 more time. Aspirate 1x Wash Buffer II.

Add Biotinylated Antibody Cocktail into each well and incubate on shaker (RT) at slow speed for 2 hours. Aspirate Biotinylated Antibody Cocktail. Add 2mL of 1x Wash Buffer I to each well and incubate on shaker (RT) at slow speed for 5 minutes. Repeat 2 more times. Aspirate 1x Wash Buffer I. Add 2mL of 1x Wash Buffer II to each well and incubate on shaker (RT) at slow speed. Repeat 1 more time. Aspirate 1x Wash Buffer II.

Add HRP-Streptavidin into each well and incubate on a shaker (RT) at a slow speed for 2 hours. Aspirate HRP-Streptavidin. Add 2mL of 1x Wash Buffer I to each well and incubate on a shaker (RT) at slow speed for 5 minutes. Repeat 2 more times. Aspirate 1x Wash Buffer I. Add 2mL of 1x Wash Buffer II to each well and incubate on a shaker (RT) slowly. Repeat 1 more time. Aspirate 1x Wash Buffer II. Pick up the membrane and remove the excess solution by touching the kimwipe. Lay on a plastic sheet. Add as many membranes as needed. Add 400µL of Detection Buffer onto each membrane and wait 2 minutes. Carefully sandwich membranes with another plastic sheet. Image.

### Western blot

For the analysis, a sample from protein fraction (separated from mitochondria) was separated on a 4-12% Bolt Bis-Tris gel (Invitrogen, NW04125BOX) and transferred using an iBlot2 Transfer system (Invitrogen) to PVDF membranes (Invitrogen, IB242002) at 23 V for 3[minutes. Membranes were blocked for 5 minutes with blocking buffer, then incubated with primary antibodies diluted in blocking buffer on a rocker 4[°C overnight. The following day, membranes were washed at room temperature in 4 10-minute washes in TBST. Membranes were incubated in a secondary antibody solution for 1 h in blocking buffer on a rocker 4[°C. Membranes were washed 4 10-minute washes in TBST at room temperature; the membranes were incubated for 5[minutes in the dark with SuperSignal™ West Pico Plus substrate ((Invitrogen, 34577) used for actin and Tuj1 detection) or SuperSignal™ West Atto Ultimate Sensitivity substrate ((Invitrogen, A38555) used for detection of the rest of the proteins) to identify the proteins of interest. The membranes were serially imaged using a BioRad ChemiDoc MP. The images were analyzed using ImageJ Software.

Western blot Primary: pDRP1 (S616) Rb (Cell Signaling 3455S), DRP1 (DNM1L) Ms (Invitrogen REF MA5-26255), TOMM20 Rb (Invitrogen REF MA5-24859), MTCO1 Ms (Invitrogen REF 459600), VDAC1 Rb (Cell Signalling D73D12), pTau (Ser202 & Thr205) Ms (Invitrogen REF MN1020), TDP43 Rb (Abcam ab109535), APP Rb (Invitrogen REF PA5-17829), Connexin 43 Rb (Cell Signal 3512), Annexin A2 Ms (Invitrogen REF 03-4400). Western Blot Secondary: Anti-rabbit IgG, HRP-linked (Cell Signaling 7074), Anti-mouse IgG, HRP-linked (Cell Signaling 7076)

### FAC Sorting for 2D transduced cell lines

Before FAC Sorting, 4 different samples must be prepared: control cells without Propidium Iodide (PI), control cells with PI, transduced cells without PI, and transduced cells with PI. Collect all MOIs of cells (including control) by aspirating media, adding 3 mL of trypsin (microglia and astrocytes) for 3 min or Trip-LE (hi-NSC) for 1 min in 37 C incubator, hit dish and add 3 mL of corresponding cell type media, collect in individual tubes. Count cells. For control cells, split evenly into two groups. For the transduced cell line with the brightest fluorescence, separate 10% of the cells to a separate tube. Centrifuge all cells at 1000 rpm for 5 min (microglia and astrocyte) or 3000 rpm for 2 min (hi-NSC). Aspirate supernatant. For all tubes, add cell sorting buffer (composed of 1x PBS, 1% Bovine Serum Albumin (ThermoFisher NC9871802), and 25 mM HEPES buffer (ThermoFisher 11-039-021)) based on 20,000,000 cells/mL. For one of the control tubes and the separated number of cells from the brightest fluorescent cell line, don’t add PI. For the rest of the tubes, add PI based on 1 microL/mL. Filter cells with 35 µm strainer cap on 5 mL polystyrene FACS tubes. The FACS machines at the Cincinnati Children’s Hospital Medical Center included BD FACSAria II.

### FACS for cell-specific mitochondria sorting

Before FAC Sorting, 9 different samples must be prepared for DNA Quantification: sham scaffolds, CCI scaffolds, NAM control scaffolds, monoculture RFP-iNSC, monoculture GFP-astrocytes, monoculture BFP-HMC3, monoculture iNSC, monoculture astrocytes, and monoculture microglia. We used Seahorse Media (50mL F12 DMEM/F-12 (Gibco 21041025), 0.5mL pyruvate, 0.5mL glucose, 0.5mL glutamine) previously mentioned to resuspend our samples. The MitoTracker Stock Solution is composed of 92µL DMSO + 50µg DeepRed MitoTracker, while the MitoTracker Working Solution is made of 0.5µL of MitoTracker stock + 250µL SH buffer. 5% Triton X: 2.5mL Triton X + 47.5mL PBS (vortex until mixed).

Thaw cryovials of cells spun down according to cell culture protocol, aspirate supernatant, and add to 2 mL tube. Use tweezers to transfer designated scaffolds to 2 mL tubes. Add 800 µL of Reagent A to all tubes. Use a P1000 pipette tip to release cells in scaffolds. Proceed with mitochondria isolation protocol. Split NAM control solution. Add 15 µL of MitoTracker working solution to all samples except one of the NAM control solutions. Take samples to the facility and proceed with sorting.

### Aspect Ratio

Mitochondrial aspect ratio (length-to-width ratio) was measured using a macro available in NIH ImageJ software, following a previously published protocol [cite here]. Maximum projection images of mitochondria, labeled with cell-specific fluorescent markers (red for neurons, blue for microglia, and green for astrocytes), were acquired using a [insert Nikon model here] confocal microscope and exported prior to analysis. The exported images were processed using the ImageJ macro to calculate the aspect ratio (AR) for each mitochondrion. Measurements were obtained from at least four images per sample, and the AR values were plotted as a distribution graph for comparative analysis across experimental groups.

### ELISA

Amyloid B 40 and Amyloid B 42 were measured using previously established protocols with slight optimization to better fit our system. A serial dilution of a prepared 2000pg/mL human amyloid beta standard was used to generate a standard curve. 50uL each of either sample or standard was added to the appropriate wells, followed by 50uL of Detection antibody. The plate was gently agitated to mix, and the incubated overnight on a rocker at 4* Celsius. After incubation, 4 instant washes were conducted using 1x wash buffer. Following the washes, 100ul of Anti-Rabbit HRP was added to each well and incubated at room temperature for 30 minutes on a shaker. Afterwards 100ul of stabilized chromogen was added, and the plate was placed in a dark environment and periodically observed to ensure over-saturation does not occur. Once optimal color development was achieved, 100ul of stop solution was added to each well. The plate was then read at 450nm, and a standard curve was applied to normalize and extract data. Data were reported as the ratio of Ab42 to Ab40.

### Messenger RNA (mRNA) sequencing and bioinformatic analysis

Approximately 1 μg of total RNA for each sample underwent RNA quality control analyses before library preparation and bulk mRNA sequencing, as previously described [50, 51]. RNA integrity numbers (RIN) were determined with a Bioanalyzer 2100 (Agilent Technologies, Santa Clara, CA, USA), which indicated minimal RNA degradation. Illumina Novaseq sequencers generated paired-end reads (150 bp) at a read depth of 20 million reads [50].

Azenta processed raw .fastq files and returned counts with featureCounts. DESeq2 R package was used to normalize and derive differentially expressed genes by comparing CCI and sham groups 6 hours and 48 hours post-injury. Comparing injury and Sham conditions, with or without P110 treatment. Gene set enrichment analyses (GSEA) were performed using the desktop software (https://www.gsea-msigdb.org/gsea/index.jsp) [56, 57] with MitoCarta3.0 [58]. Heatmaps were generated using ggplot and Illustrator. Gene module co-expression analyses were performed with the CEMiTool R package [60, 61], with the Reactome pathway database (https://reactome.org/), and signed interaction network analysis utilizing high confidence STRING protein-protein interactions (STRING score > 700). Oxidative phosphorylation complex gene set used [59] dataset.

## References

1 Dewan, M. C. et al. Estimating the global incidence of traumatic brain injury. J Neurosurg, 1–18 (2018). 10.3171/2017.10.JNS17352

2 Osier, N. D., Carlson, S. W., DeSana, A. & Dixon, C. E. Chronic Histopathological and Behavioral Outcomes of Experimental Traumatic Brain Injury in Adult Male Animals. J Neurotrauma 32, 1861–1882 (2015). 10.1089/neu.2014.3680

3 Villapol, S., Byrnes, K. R. & Symes, A. J. Temporal dynamics of cerebral blood flow, cortical damage, apoptosis, astrocyte-vasculature interaction and astrogliosis in the pericontusional region after traumatic brain injury. Front Neurol 5, 82 (2014). 10.3389/fneur.2014.00082

4 Ladak, A. A., Enam, S. A. & Ibrahim, M. T. A Review of the Molecular Mechanisms of Traumatic Brain Injury. World Neurosurg 131, 126–132 (2019). 10.1016/j.wneu.2019.07.039

5 Johnson, V. E., Stewart, W. & Smith, D. H. Traumatic brain injury and amyloid-beta pathology: a link to Alzheimer’s disease? Nat Rev Neurosci 11, 361–370 (2010). 10.1038/nrn2808

6 Ramos-Cejudo, J. et al. Traumatic Brain Injury and Alzheimer’s Disease: The Cerebrovascular Link. EBioMedicine 28, 21–30 (2018). 10.1016/j.ebiom.2018.01.021

7 Wu, D. et al. Traumatic Brain Injury Accelerates the Onset of Cognitive Dysfunction and Aggravates Alzheimer’s-Like Pathology in the Hippocampus by Altering the Phenotype of Microglia in the APP/PS1 Mouse Model. Front Neurol 12, 666430 (2021). 10.3389/fneur.2021.666430

8 Srinivasan, G. & Brafman, D. A. The Emergence of Model Systems to Investigate the Link Between Traumatic Brain Injury and Alzheimer’s Disease. Front Aging Neurosci 13, 813544 (2021). 10.3389/fnagi.2021.813544

9 Cherry, J. D. et al. Microglial neuroinflammation contributes to tau accumulation in chronic traumatic encephalopathy. Acta Neuropathol Commun 4, 112 (2016). 10.1186/s40478-016-0382-8

10 Bader, V. & Winklhofer, K. F. Mitochondria at the interface between neurodegeneration and neuroinflammation. Semin Cell Dev Biol 99, 163–171 (2020). 10.1016/j.semcdb.2019.05.028

11 Herst, P. M., Rowe, M. R., Carson, G. M. & Berridge, M. V. Functional Mitochondria in Health and Disease. Front Endocrinol (Lausanne) 8, 296 (2017). 10.3389/fendo.2017.00296

12 Norat, P. et al. Mitochondrial dysfunction in neurological disorders: Exploring mitochondrial transplantation. NPJ Regen Med 5, 22 (2020). 10.1038/s41536-020-00107-x

13 Xing, G., Ren, M., Watson, W. D., O’Neill, J. T. & Verma, A. Traumatic brain injury-induced expression and phosphorylation of pyruvate dehydrogenase: a mechanism of dysregulated glucose metabolism. Neurosci Lett 454, 38–42 (2009). 10.1016/j.neulet.2009.01.047

14 Huynh, L. M., Burns, M. P., Taub, D. D., Blackman, M. R. & Zhou, J. Chronic Neurobehavioral Impairments and Decreased Hippocampal Expression of Genes Important for Brain Glucose Utilization in a Mouse Model of Mild TBI. Front Endocrinol (Lausanne) 11, 556380 (2020). 10.3389/fendo.2020.556380

15 Xing, G. et al. Pyruvate dehydrogenase phosphatase1 mRNA expression is divergently and dynamically regulated between rat cerebral cortex, hippocampus and thalamus after traumatic brain injury: a potential biomarker of TBI-induced hyper- and hypo- glycaemia and neuronal vulnerability. Neurosci Lett 525, 140–145 (2012). 10.1016/j.neulet.2012.07.055

16 Hubbard, W. B., Harwood, C. L., Geisler, J. G., Vekaria, H. J. & Sullivan, P. G. Mitochondrial uncoupling prodrug improves tissue sparing, cognitive outcome, and mitochondrial bioenergetics after traumatic brain injury in male mice. J Neurosci Res 96, 1677–1688 (2018). 10.1002/jnr.24271

17 Hubbard, W. B. et al. Acute Mitochondrial Impairment Underlies Prolonged Cellular Dysfunction after Repeated Mild Traumatic Brain Injuries. J Neurotrauma 36, 1252–1263 (2019). 10.1089/neu.2018.5990

18 Lyons, D. N. et al. A Mild Traumatic Brain Injury in Mice Produces Lasting Deficits in Brain Metabolism. J Neurotrauma 35, 2435–2447 (2018). 10.1089/neu.2018.5663

19 Yonutas, H. M., Vekaria, H. J. & Sullivan, P. G. Mitochondrial specific therapeutic targets following brain injury. Brain Res 1640, 77–93 (2016). 10.1016/j.brainres.2016.02.007

20 Arun, P. et al. Acute mitochondrial dysfunction after blast exposure: potential role of mitochondrial glutamate oxaloacetate transaminase. J Neurotrauma 30, 1645–1651 (2013). 10.1089/neu.2012.2834

21 Agrawal, R. et al. Dietary fructose aggravates the pathobiology of traumatic brain injury by influencing energy homeostasis and plasticity. J Cereb Blood Flow Metab 36, 941–953 (2016). 10.1177/0271678X15606719

22 Hill, R. L., Singh, I. N., Wang, J. A., Kulbe, J. R. & Hall, E. D. Protective effects of phenelzine administration on synaptic and non-synaptic cortical mitochondrial function and lipid peroxidation-mediated oxidative damage following TBI in young adult male rats. Exp Neurol 330, 113322 (2020). 10.1016/j.expneurol.2020.113322

23 Lamade, A. M., et al. Mitochondrial damage & lipid signaling in traumatic brain injury. Exp Neurol 329, 113307 (2020). 10.1016/j.expneurol.2020.113307

24 Vike, N. L. et al. A preliminary model of football-related neural stress that integrates metabolomics with transcriptomics and virtual reality. iScience 25, 103483 (2022). 10.1016/j.isci.2021.103483

25 Bari, S., et al. Integrating multi-omics with neuroimaging and behavior: A preliminary model of dysfunction in football athletes. Neuroimage: Reports 1 (2021). 10.1016/j.ynirp.2021.100032

26 Bhatti, J. S., Bhatti, G. K. & Reddy, P. H. Mitochondrial dysfunction and oxidative stress in metabolic disorders - A step towards mitochondria based therapeutic strategies. Biochim Biophys Acta Mol Basis Dis 1863, 1066–1077 (2017). 10.1016/j.bbadis.2016.11.010

27 Ryu, W. I. et al. Brain cells derived from Alzheimer’s disease patients have multiple specific innate abnormalities in energy metabolism. Mol Psychiatry 26, 5702–5714 (2021). 10.1038/s41380-021-01068-3

28 Yu, L., Jin, J., Xu, Y. & Zhu, X. Aberrant Energy Metabolism in Alzheimer’s Disease. J Transl Int Med 10, 197–206 (2022). 10.2478/jtim-2022-0024

29 Liaudanskaya, V. et al. Modeling Controlled Cortical Impact Injury in 3D Brain-Like Tissue Cultures. Adv Healthc Mater 9, e2000122 (2020). 10.1002/adhm.202000122

30 Liaudanskaya, V. et al. Mitochondria dysregulation contributes to secondary neurodegeneration progression post-contusion injury in human 3D in vitro triculture brain tissue model. Cell Death Dis 14, 496 (2023). 10.1038/s41419-023-05980-0

31 Rouleau, N. et al. A Long-Living Bioengineered Neural Tissue Platform to Study Neurodegeneration. Macromol Biosci 20, e2000004 (2020). 10.1002/mabi.202000004

32 Snapper, D. M. et al. Development of a novel bioengineered 3D brain-like tissue for studying primary blast-induced traumatic brain injury. J Neurosci Res 101, 3–19 (2023). 10.1002/jnr.25123

33 Tang-Schomer, M. D. et al. Bioengineered functional brain-like cortical tissue. Proc Natl Acad Sci U S A 111, 13811–13816 (2014). 10.1073/pnas.1324214111

34 Chiaradia, I. & Lancaster, M. A. Brain organoids for the study of human neurobiology at the interface of in vitro and in vivo. Nat Neurosci 23, 1496–1508 (2020). 10.1038/s41593-020-00730-3

35 Lancaster, M. A. et al. Cerebral organoids model human brain development and microcephaly. Nature 501, 373–379 (2013). 10.1038/nature12517

36 Garza, R. et al. Single-cell transcriptomics of human traumatic brain injury reveals activation of endogenous retroviruses in oligodendroglia. Cell Rep 42, 113395 (2023). 10.1016/j.celrep.2023.113395

37 Russo, P. S. T. et al. CEMiTool: a Bioconductor package for performing comprehensive modular co-expression analyses. BMC Bioinformatics 19, 56 (2018). 10.1186/s12859-018-2053-1

38 Langfelder, P. & Horvath, S. WGCNA: an R package for weighted correlation network analysis. BMC Bioinformatics 9, 559 (2008). 10.1186/1471-2105-9-559

39 Subramanian, A. et al. Gene set enrichment analysis: a knowledge-based approach for interpreting genome-wide expression profiles. Proc Natl Acad Sci U S A 102, 15545–15550 (2005). 10.1073/pnas.0506580102

40 Mootha, V. K. et al. PGC-1alpha-responsive genes involved in oxidative phosphorylation are coordinately downregulated in human diabetes. Nat Genet 34, 267–273 (2003). 10.1038/ng1180

41 Rath, S. et al. MitoCarta3.0: an updated mitochondrial proteome now with sub-organelle localization and pathway annotations. Nucleic Acids Res 49, D1541–D1547 (2021). 10.1093/nar/gkaa1011

42 Guarnieri, J. W. et al. Core mitochondrial genes are down-regulated during SARS-CoV-2 infection of rodent and human hosts. Sci Transl Med 15, eabq1533 (2023). 10.1126/scitranslmed.abq1533

43 Abu Hamdeh, S., et al. Differential DNA Methylation of the Genes for Amyloid Precursor Protein, Tau, and Neurofilaments in Human Traumatic Brain Injury. J Neurotrauma 38, 1679–1688 (2021). 10.1089/neu.2020.7283

44 Sabbir, M. G. Loss of calcium/calmodulin-dependent protein kinase kinase 2, transferrin, and transferrin receptor proteins in the temporal cortex of Alzheimer’s patients postmortem is associated with abnormal iron homeostasis: implications for patient survival. Front Cell Dev Biol 12, 1469751 (2024). 10.3389/fcell.2024.1469751

45 Ghosh, A. & Giese, K. P. Calcium/calmodulin-dependent kinase II and Alzheimer’s disease. Mol Brain 8, 78 (2015). 10.1186/s13041-015-0166-2

46 Abiose, O. et al. Post-translational modifications linked to preclinical Alzheimer’s disease-related pathological and cognitive changes. Alzheimers Dement 20, 1851–1867 (2024). 10.1002/alz.13576

47 Jaisa-Aad, M., Munoz-Castro, C., Healey, M. A., Hyman, B. T. & Serrano-Pozo, A. Characterization of monoamine oxidase-B (MAO-B) as a biomarker of reactive astrogliosis in Alzheimer’s disease and related dementias. Acta Neuropathol 147, 66 (2024). 10.1007/s00401-024-02712-2

48 Tang, X. et al. Research progress on SLC7A11 in the regulation of cystine/cysteine metabolism in tumors. Oncol Lett 23, 47 (2022). 10.3892/ol.2021.13165

49 Escartin, C. et al. Reactive astrocyte nomenclature, definitions, and future directions. Nat Neurosci 24, 312–325 (2021). 10.1038/s41593-020-00783-4

50 Wei, Z., Koya, J. & Reznik, S. E. Insulin Resistance Exacerbates Alzheimer Disease via Multiple Mechanisms. Front Neurosci 15, 687157 (2021). 10.3389/fnins.2021.687157

51 Gonzalez-Rodriguez, M. et al. Neurodegeneration and Astrogliosis in the Human CA1 Hippocampal Subfield Are Related to hsp90ab1 and bag3 in Alzheimer’s Disease. Int J Mol Sci 23 (2021). 10.3390/ijms23010165

52 da Costa, I. B., et al. Change in INSR, APBA2 and IDE Gene Expressions in Brains of Alzheimer’s Disease Patients. Curr Alzheimer Res 14, 760–765 (2017). 10.2174/1567205014666170203100734

53 Folch, J. et al. The Implication of the Brain Insulin Receptor in Late Onset Alzheimer’s Disease Dementia. Pharmaceuticals (Basel) 11 (2018). 10.3390/ph11010011

54 Lackie, R. E. et al. Increased levels of Stress-inducible phosphoprotein-1 accelerates amyloid-beta deposition in a mouse model of Alzheimer’s disease. Acta Neuropathol Commun 8, 143 (2020). 10.1186/s40478-020-01013-5

55 Zhang, B. et al. Integrated systems approach identifies genetic nodes and networks in late-onset Alzheimer’s disease. Cell 153, 707–720 (2013). 10.1016/j.cell.2013.03.030

56 Cermak, S. et al. Loss of Cathepsin B and L Leads to Lysosomal Dysfunction, NPC-Like Cholesterol Sequestration and Accumulation of the Key Alzheimer’s Proteins. PLoS One 11, e0167428 (2016). 10.1371/journal.pone.0167428

57 Kim, A. & Benavente, C. A. Oncogenic Roles of UHRF1 in Cancer. Epigenomes 8 (2024). 10.3390/epigenomes8030026

58 Miyashita, R. et al. The termination of UHRF1-dependent PAF15 ubiquitin signaling is regulated by USP7 and ATAD5. Elife 12 (2023). 10.7554/eLife.79013

59 Wei-Shan, H., Amit, V. C. & Clarke, D. J. Cell cycle regulation of condensin Smc4. Oncotarget 10, 263–276 (2019). 10.18632/oncotarget.26467

60 Ramachandran, K. et al. Transcriptional programming of translation by BCL6 controls skeletal muscle proteostasis. Nat Metab 6, 304–322 (2024). 10.1038/s42255-024-00983-3

61 Baskin, K. K. et al. MED12 regulates a transcriptional network of calcium-handling genes in the heart. JCI Insight 2 (2017). 10.1172/jci.insight.91920

62 Dabovic, B. et al. Control of lung development by latent TGF-beta binding proteins. J Cell Physiol 226, 1499–1509 (2011). 10.1002/jcp.22479

63 Moon, D. O. Deciphering the Role of BCAR3 in Cancer Progression: Gene Regulation, Signal Transduction, and Therapeutic Implications. Cancers (Basel) 16 (2024). 10.3390/cancers16091674

64 Taiko, I. et al. Selection of red fluorescent protein for genetic labeling of mitochondria and intercellular transfer of viable mitochondria. Sci Rep 12, 19841 (2022). 10.1038/s41598-022-24297-0

65 Carpenter, K. L., Jalloh, I. & Hutchinson, P. J. Glycolysis and the significance of lactate in traumatic brain injury. Front Neurosci 9, 112 (2015). 10.3389/fnins.2015.00112

66 Stovell, M. G. et al. Assessing Metabolism and Injury in Acute Human Traumatic Brain Injury with Magnetic Resonance Spectroscopy: Current and Future Applications. Front Neurol 8, 426 (2017). 10.3389/fneur.2017.00426

67 Xu, X. J. et al. Glucose metabolism: A link between traumatic brain injury and Alzheimer’s disease. Chin J Traumatol 24, 5–10 (2021). 10.1016/j.cjtee.2020.10.001

68 Kajiwara, Y. et al. GJA1 (connexin43) is a key regulator of Alzheimer’s disease pathogenesis. Acta Neuropathol Commun 6, 144 (2018). 10.1186/s40478-018-0642-x

69 Maulik, M. et al. Amyloid-beta regulates gap junction protein connexin 43 trafficking in cultured primary astrocytes. J Biol Chem 295, 15097–15111 (2020). 10.1074/jbc.RA120.013705

70 White, Z. B., 2nd, Nair, S. & Bredel, M. The role of annexins in central nervous system development and disease. J Mol Med (Berl) 102, 751–760 (2024). 10.1007/s00109-024-02443-7

71 Gauthier-Kemper, A. et al. Annexins A2 and A6 interact with the extreme N terminus of tau and thereby contribute to tau’s axonal localization. J Biol Chem 293, 8065–8076 (2018). 10.1074/jbc.RA117.000490

72 Astillero-Lopez, V. et al. Proteomic analysis identifies HSP90AA1, PTK2B, and ANXA2 in the human entorhinal cortex in Alzheimer’s disease: Potential role in synaptic homeostasis and Abeta pathology through microglial and astroglial cells. Brain Pathol 34, e13235 (2024). 10.1111/bpa.13235

73 Lauro, C. & Limatola, C. Metabolic Reprograming of Microglia in the Regulation of the Innate Inflammatory Response. Front Immunol 11, 493 (2020). 10.3389/fimmu.2020.00493

74 Strope, T. A., Birky, C. J. & Wilkins, H. M. The Role of Bioenergetics in Neurodegeneration. Int J Mol Sci 23 (2022). 10.3390/ijms23169212

75 Zong, Y. et al. Mitochondrial dysfunction: mechanisms and advances in therapy. Signal Transduct Target Ther 9, 124 (2024). 10.1038/s41392-024-01839-8

76 Cleland, N. R. W., Al-Juboori, S. I., Dobrinskikh, E. & Bruce, K. D. Altered substrate metabolism in neurodegenerative disease: new insights from metabolic imaging. J Neuroinflammation 18, 248 (2021). 10.1186/s12974-021-02305-w

77 Sundman, M. H., Hall, E. E. & Chen, N. K. Examining the relationship between head trauma and neurodegenerative disease: A review of epidemiology, pathology and neuroimaging techniques. J Alzheimers Dis Parkinsonism 4 (2014). 10.4172/2161-0460.1000137

78 Picard, M. & Shirihai, O. S. Mitochondrial signal transduction. Cell Metab 34, 1620–1653 (2022). 10.1016/j.cmet.2022.10.008

79 Gardner, R. C. et al. Mild TBI and risk of Parkinson’s disease: A chronic effect of Neurotrauma Consortium Study. Neurology 90 (2018).

80 Lizhnyak, P. N. & Ottens, A. K. Proteomics: in pursuit of effective traumatic brain injury therapeutics. Expert Review of Proteomics 12 (2015).

81 Borst, K., Schwabenland, M. & Prinz, M. Microglia metabolism in health and disease. Neurochem Int 130, 104331 (2019). 10.1016/j.neuint.2018.11.006

82 Brett, B. L., Gardner, R. C., Godbout, J., Dams-O’Connor, K. & Keene, C. D. Traumatic Brain Injury and Risk of Neurodegenerative Disorder. Biol Psychiatry 91, 498–507 (2022). 10.1016/j.biopsych.2021.05.025

83 Guerrero, A., De Strooper, B. & Arancibia-Carcamo, I. L. Cellular senescence at the crossroads of inflammation and Alzheimer’s disease. Trends Neurosci 44, 714–727 (2021). 10.1016/j.tins.2021.06.007

84 Haukedal, H. & Freude, K. K. Implications of Glycosylation in Alzheimer’s Disease. Front Neurosci 14, 625348 (2020). 10.3389/fnins.2020.625348

85 Lin, A. P., et al. MYC, mitochondrial metabolism and O-GlcNAcylation converge to modulate the activity and subcellular localization of DNA and RNA demethylases. Leukemia 36, 1150–1159 (2022). 10.1038/s41375-021-01489-7

86 Tracey, T. J., Steyn, F. J., Wolvetang, E. J. & Ngo, S. T. Neuronal Lipid Metabolism: Multiple Pathways Driving Functional Outcomes in Health and Disease. Front Mol Neurosci 11, 10 (2018). 10.3389/fnmol.2018.00010

87 Chakraborty, R., Nonaka, T., Hasegawa, M. & Zurzolo, C. Tunnelling nanotubes between neuronal and microglial cells allow bi-directional transfer of alpha-Synuclein and mitochondria. Cell Death Dis 14, 329 (2023). 10.1038/s41419-023-05835-8

88 Cheng, Y. & Bai, F. The Association of Tau With Mitochondrial Dysfunction in Alzheimer’s Disease. Front Neurosci 12, 163 (2018). 10.3389/fnins.2018.00163

89 Pan, K. H., Chang, H. & Yang, W. Y. Extracellular release in the quality control of the mammalian mitochondria. J Biomed Sci 30, 85 (2023). 10.1186/s12929-023-00979-3

90 Pinto, G. et al. Patient-derived glioblastoma stem cells transfer mitochondria through tunneling nanotubes in tumor organoids. Biochem J 478, 21–39 (2021). 10.1042/BCJ20200710

91 Scheiblich, H. et al. Microglia jointly degrade fibrillar alpha-synuclein cargo by distribution through tunneling nanotubes. Cell 184, 5089–5106 e5021 (2021). 10.1016/j.cell.2021.09.007

92 Zampieri, L. X., Silva-Almeida, C., Rondeau, J. D. & Sonveaux, P. Mitochondrial Transfer in Cancer: A Comprehensive Review. Int J Mol Sci 22 (2021). 10.3390/ijms22063245

93 Zhao, Z. et al. Extracellular mitochondria released from traumatized brains induced platelet procoagulant activity. Haematologica 105, 209–217 (2020). 10.3324/haematol.2018.214932

94 Zhang, Y. et al. Synergistic label-free fluorescence imaging and miRNA studies reveal dynamic human neuron-glial metabolic interactions following injury. Sci Adv 10, eadp1980 (2024). 10.1126/sciadv.adp1980

95 Hubbard, W. B. et al. Mitochondrial Dysfunction After Repeated Mild Blast Traumatic Brain Injury Is Attenuated by a Mild Mitochondrial Uncoupling Prodrug. J Neurotrauma 40, 2396–2409 (2023). 10.1089/neu.2023.0102

96 Vafai, S. B. & Mootha, V. K. Mitochondrial disorders as windows into an ancient organelle. Nature 491, 374–383 (2012). 10.1038/nature11707

97 Mattson, M. P., Gleichmann, M. & Cheng, A. Mitochondria in neuroplasticity and neurological disorders. Neuron 60, 748–766 (2008). 10.1016/j.neuron.2008.10.010

98 Cherry, J. D., Olschowka, J. A. & O’Banion, M. K. Neuroinflammation and M2 microglia: the good, the bad, and the inflamed. J Neuroinflammation 11, 98 (2014). 10.1186/1742-2094-11-98

99 Cheng, A., Hou, Y. & Mattson, M. P. Mitochondria and neuroplasticity. ASN Neuro 2, e00045 (2010). 10.1042/AN20100019

100 Lee, A. et al. Abeta42 oligomers trigger synaptic loss through CAMKK2-AMPK-dependent effectors coordinating mitochondrial fission and mitophagy. Nat Commun 13, 4444 (2022). 10.1038/s41467-022-32130-5

101 Devanney, N. A., Stewart, A. N. & Gensel, J. C. Microglia and macrophage metabolism in CNS injury and disease: The role of immunometabolism in neurodegeneration and neurotrauma. Exp Neurol 329, 113310 (2020). 10.1016/j.expneurol.2020.113310

102 Ashton, N. J. et al. Differential roles of Abeta42/40, p-tau231 and p-tau217 for Alzheimer’s trial selection and disease monitoring. Nat Med 28, 2555–2562 (2022). 10.1038/s41591-022-02074-w

103 Karran, E. & De Strooper, B. The amyloid hypothesis in Alzheimer disease: new insights from new therapeutics. Nat Rev Drug Discov 21, 306–318 (2022). 10.1038/s41573-022-00391-w

104 Rockwood, D. N. et al. Materials fabrication from Bombyx mori silk fibroin. Nat Protoc 6, 1612–1631 (2011). 10.1038/nprot.2011.379

105 Dingle, Y. L., Bonzanni, M., Liaudanskaya, V., Nieland, T. J. F. & Kaplan, D. L. Integrated functional neuronal network analysis of 3D silk-collagen scaffold-based mouse cortical culture. STAR Protoc 2, 100292 (2021). 10.1016/j.xpro.2020.100292

106 Cairns, D. M. et al. Expandable and Rapidly Differentiating Human Induced Neural Stem Cell Lines for Multiple Tissue Engineering Applications. Stem Cell Reports 7, 557–570 (2016). 10.1016/j.stemcr.2016.07.017

