## Supplementary Figures for "The Silent Saboteur: How Mitochondria Shape the Long-Term Fate of the Injured Brain": Kansakar_Cell Metabolism_Supplementary Document.docx

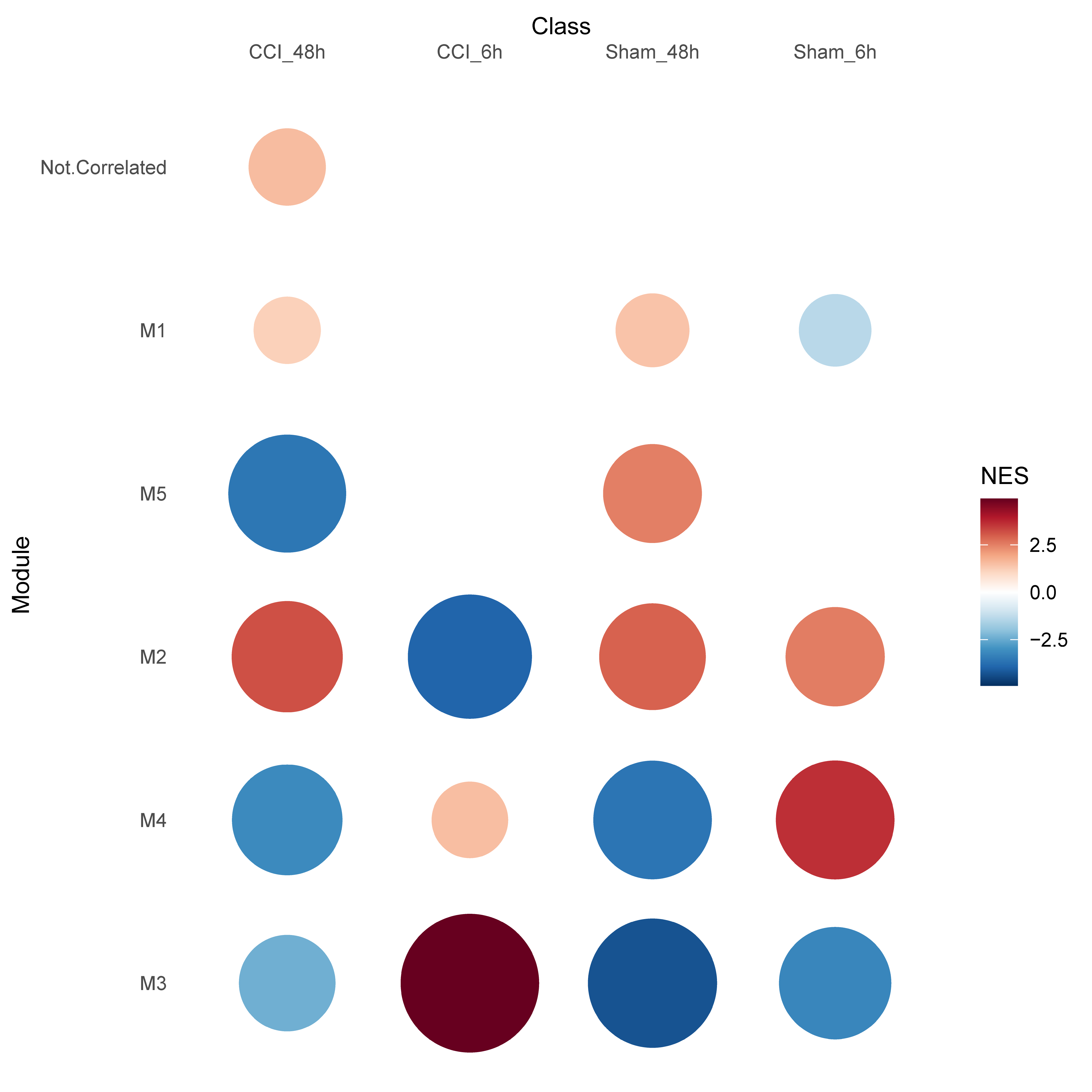

**Figure S1. Co-expression network analysis revealed a significant shift in gene expression** associated with M1 – mitochondria function (positively associated with an injury); M2 – cell cycle progression (negatively associated at 6 hours); M3 – cellular signaling (positively associated with injury at 6 hours and negatively at 48 hours); M4 – tissue development (no significant differences between sham and injury groups); M5 – neuroinflammation (negatively associated at 48 hours) in human 3D triculture model in response to contusion injury. Changes in differentially expressed genes 6 and 48 hours post-moderate injury in human 3D triculture model composed of mtDSred2-neurons, mtEGFP-astrocytes, and BFP2-microglia. Sham (n = 5) and CCI (n = 6) at 6 hours, and Sham (n = 4) and CCI (n = 4) were used to obtain data for this transcriptomic profile.

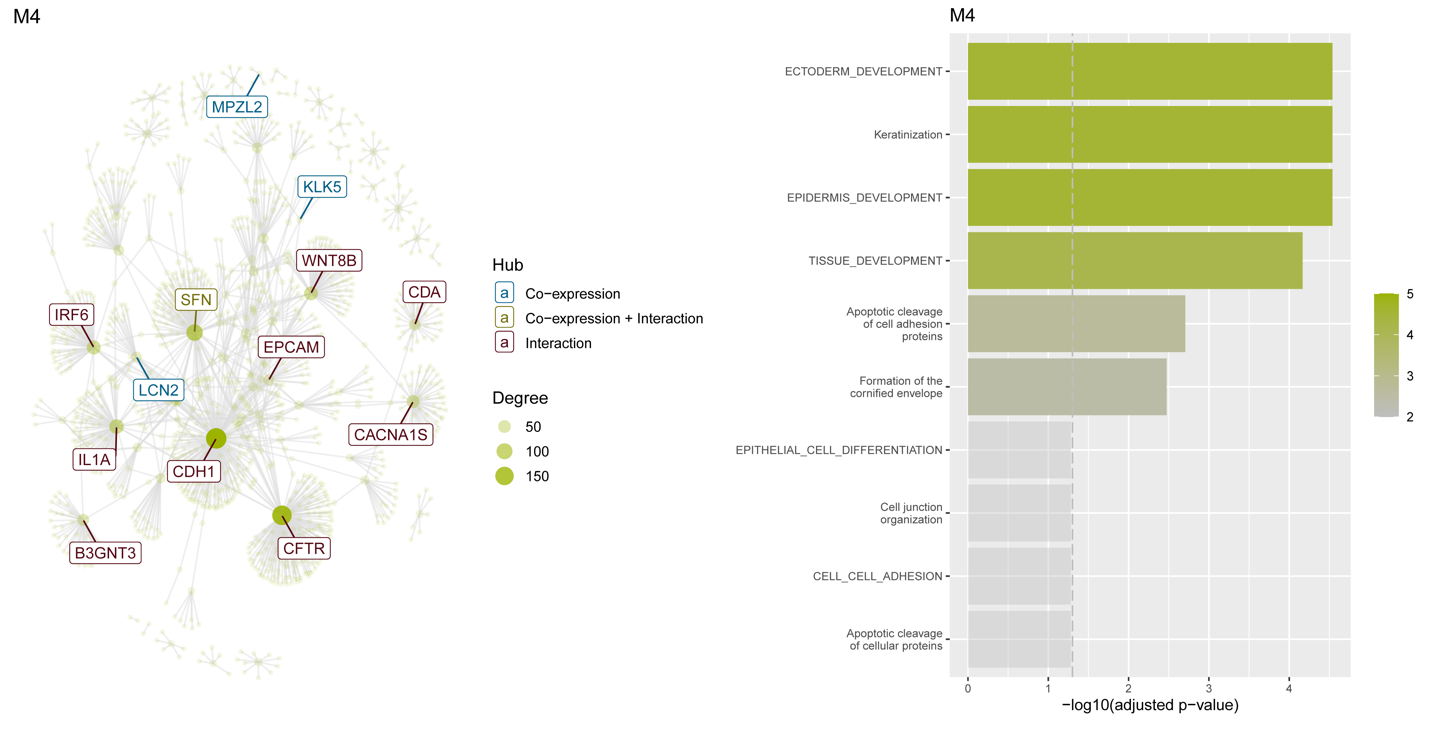

**Figure S2. Moderate injury-induced transcriptomic changes in tissue development markers** in human 3D *in vitro* brain tissues 6 and 48 hours after moderate injury inflicted by the controlled cortical impactor. The human 3D *in vitro* triculture model comprised 2 mln of neurons, 0.5 mln astrocytes, and 0.1 mln of HMC3 microglial cells. Sham (n = 5) and CCI (n = 6) at 6 hours, and Sham (n = 4) and CCI (n = 4) were used to obtain data for this transcriptomic profile. Gene set enrichment analysis revealed no significant change in Module 4 between sham and injury (involved in tissue development); however, there was a significant difference between sham and injury at 6 hours vs 48 hours.

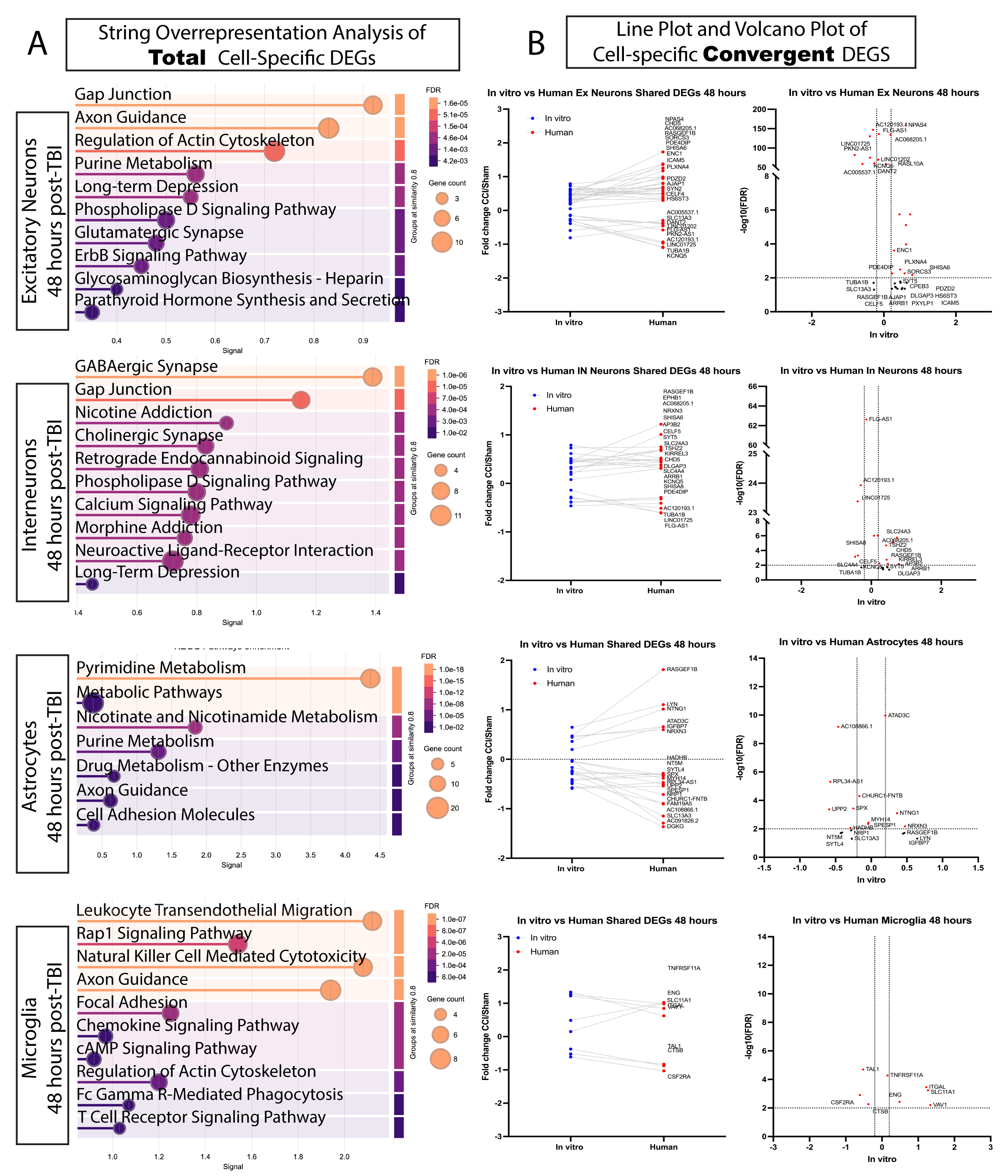

**Figure S3. Human TBI transcriptomics align with our 3D in vitro model, revealing conserved mitochondrial dysfunction.** 48-hour results supplementary to the main Figure 2. n=5 Shams and n=6 CCI samples at 6 hours and n=5 Shams and n=6 CCI samples were used to obtain data for this transcriptomic profile. Single-cell RNA sequencing from 12 critically injured or deceased TBI patients was analyzed to curate cell-specific differentially expressed genes (DEGs) across excitatory neurons, interneurons, astrocytes, and microglia, using data from five uninjured controls as a reference. This patient-derived transcriptomic dataset was then integrated with bulk RNA sequencing from human 3D in vitro brain injury samples at 6 and 48 hours post-injury. This enables a comparative analysis of conserved molecular signatures between clinical and experimental models. Differentially expressed genes (abs(Log2(Fold Change))>0.585 and q<0.01).

(A) Regardless of convergence or divergence, similar DEGs across the datasets were run with STRING analysis to determine cell-specific pathways significantly altered in response to trauma.
(B) Convergent upregulated and downregulated DEGs were isolated and plotted for 4 distinct cell populations.

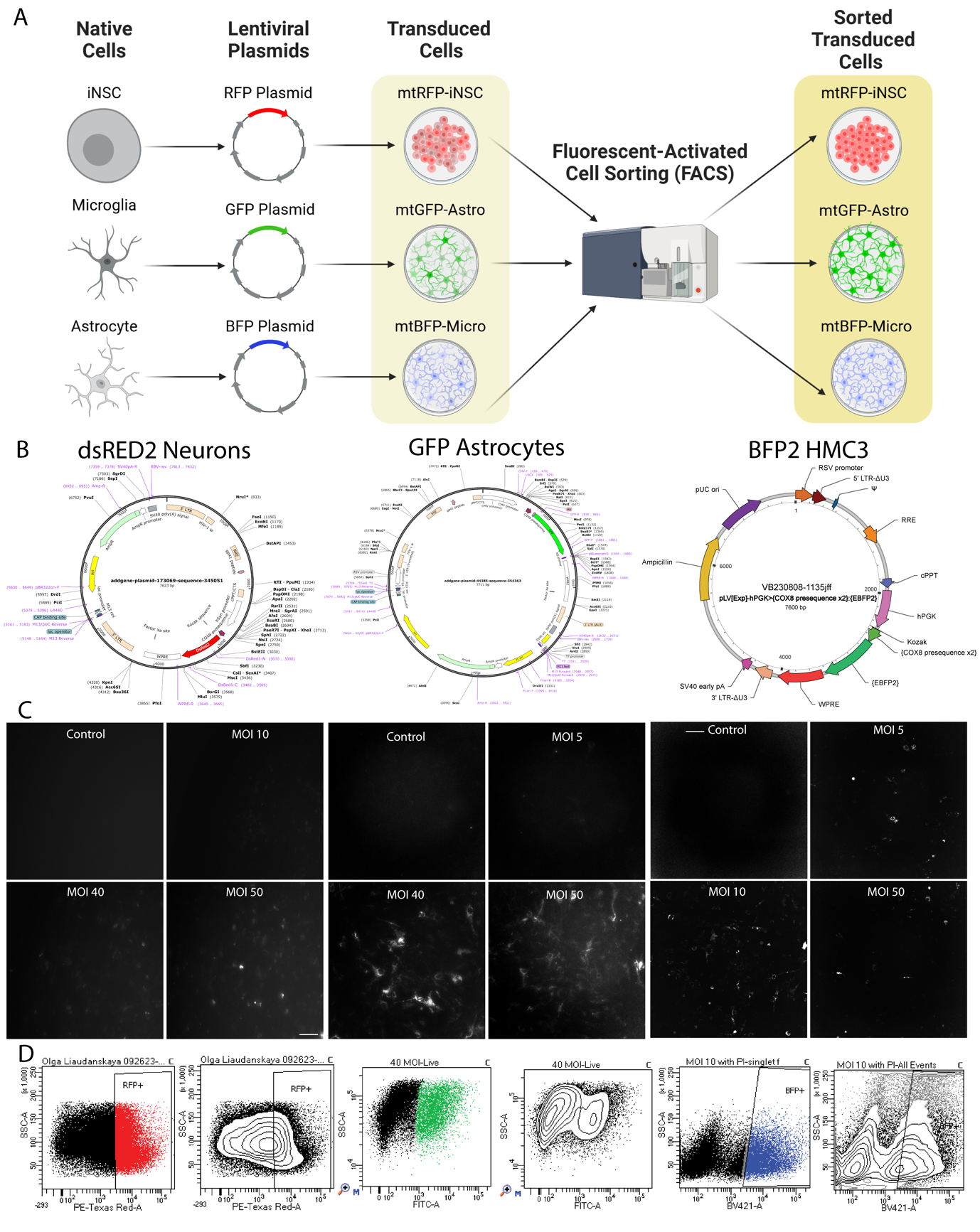

**Figure S4.** **Design and validation of fluorescently labeled mitochondria.**

(A) Schematic representation of mitochondria transduction experiments.
(B) dsRED2 Addgene plasmid used to transduce human-induced neural stem cells, EGFP Addgene plasmid used to transduce human primary astrocytes, and custom-designed BFP2 plasmid used to transduce HMC3 microglia cell line.
(C) Epifluorescent images of experimental multiplicities of infection (MOIs) in transduced fluorescent cell lines cultured in 2D, captured using a Cytation epifluorescent microscope. Scale bar: 100 μm.
(D) Fluorescent-activated cell sorter gating parameters for transduced cells with final MOIs.

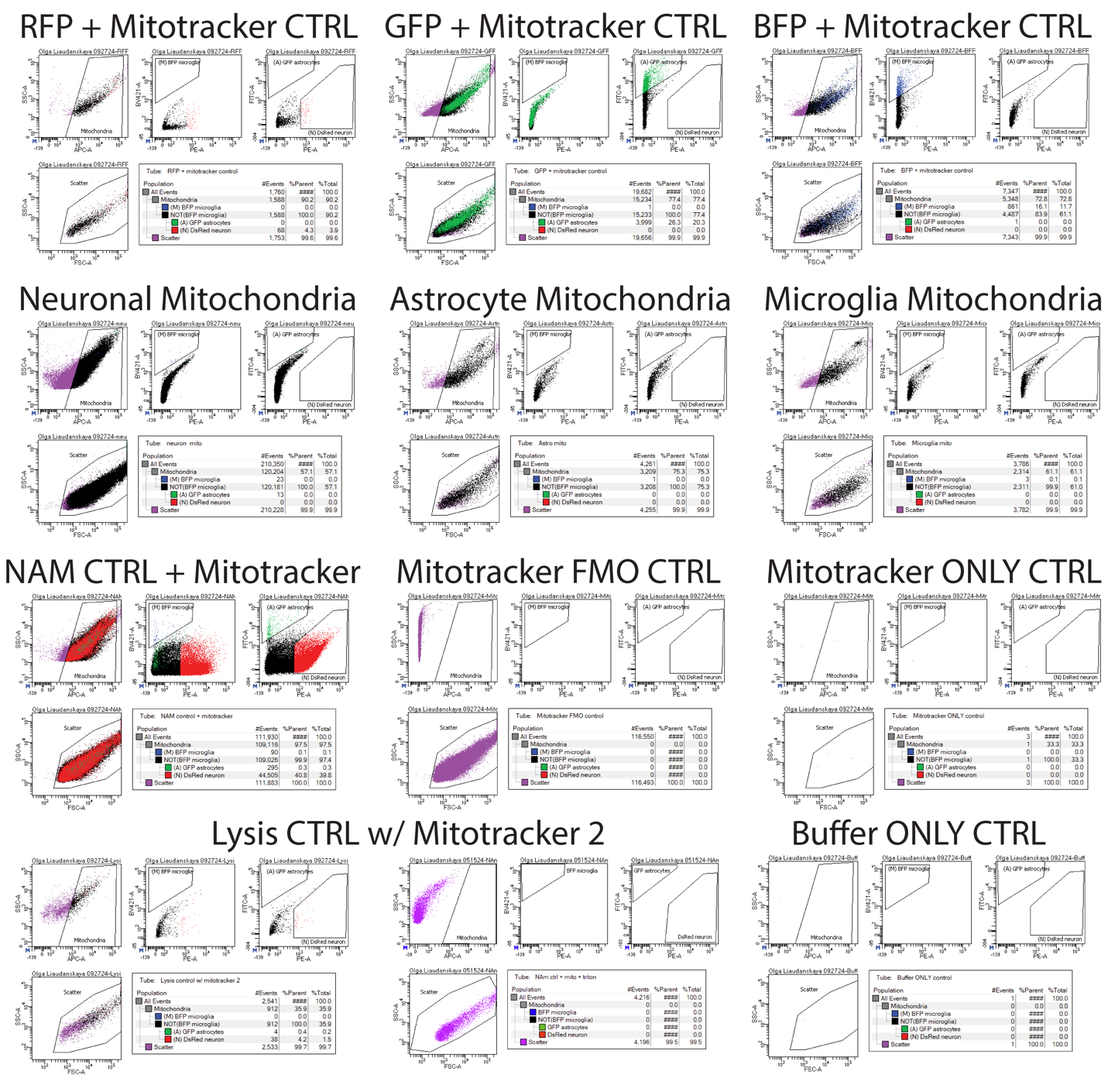

**Figure S5. Fluorescently Activated Cell Sorting of cell-specific mitochondria experimental set-up (representative panels from 1 sorting experiment).** To ensure proper mitochondria sorting with 3 different fluorophores, we used 11 control conditions for each mitochondria sorting experiment. (Top) RFP+ mitochondria from Neurons, GFP+ mitochondria from Astrocytes and BFP+ mitochondria from Microglia monocultures counterstained with MitoTracker with 647 nm wavelength; (Row 2) Neuronal, Astrocytic and Microglia mitochondria from naïve untagged monocultures but stained with 647 nm MitoTracker; (Row 3) Mitochondria isolated from NAM tricultures with fluorescently tagged mitochondria + MitoTracker in 647 nm wavelength; Mitochondria isolated from NAM tricultures with fluorescently tagged mitochondria **Without** MitoTracker in 647 nm wavelength; and Mitotracker solution only (without mitochondria); (Bottom) Lysis CTRL (Tryton X treatment) with Mitotracker from 2 sorts and FACS sorting Buffer only.

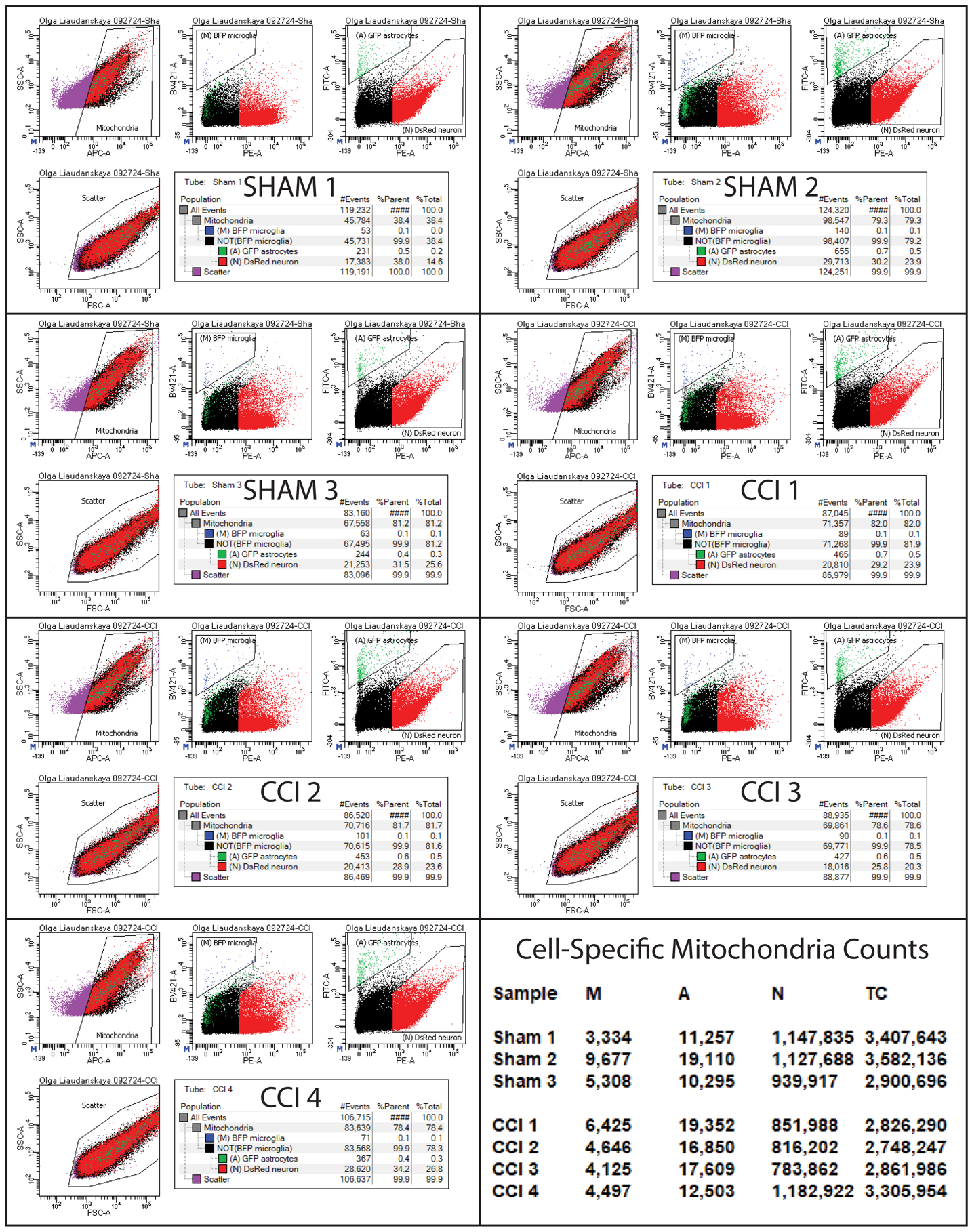

**Figure S6. Fluorescently Activated Cell Sorting of cell-specific mitochondria (representative panels from 1 sorting experiment).** Representative panels demonstrate recorded results from a one-time point sorting experiment from one biological replicate to demonstrate the gating used to separate 3 distinct mitochondria populations based on the BFP2, EGFP, and dsRED2 fluorescent tags attached.
